# Beyond Lipschitz: Ranking Binding Affinity in Hyperbolic Space

**DOI:** 10.64898/2026.02.01.703164

**Authors:** Kelin Wu, Xin Hong, Wenyu Zhu, Bowen Gao, Wei-Ying Ma, Yanyan Lan

**Affiliations:** Institute for AI Industry Research (AIR), Tsinghua University; Qiuzhen College, Tsinghua University; Department of Computer Science and Technology, Tsinghua University

## Abstract

Despite the importance of protein-ligand affinity ranking in drug discovery, existing deep learning models struggle to distinguish hard inactives that are structurally similar to active compounds but biologically inactive. We theoretically show this failure stems from the Lipschitz continuity constraint in Euclidean space, which makes neural networks locally insensitive to subtle yet critical structural perturbations. To overcome this, we propose AlphaRank, which demonstrates that scoring affinity as the negative hyperbolic geodesic distance allows subtle tangential variations to bypass Lipschitz constraints, intuitively corresponding to pivotal binding mode factors such as conformational fit. Meanwhile, AlphaRank employs joint optimization to simultaneously ranking affinity scores among actives in a proximity-aware manner and separate actives from decoy inactives through a geometric cone constraint. Experiments demonstrate that AlphaRank outperforms state-of-the-art models like Boltz-2 in both affinity ranking and active-inactive discrimination, while providing a interpretable representation space.

## 1. Introduction

Accurately ranking protein-ligand binding affinity remains a fundamental challenge in drug discovery, providing critical guidance for downstream pipelines such as lead identification and optimization. Targeting a given protein, this task primarily focuses on predicting whether a molecule is active and, if so, the extent of their binding affinity relative to other actives. Traditional physical approaches in this field have inherent limitations. For instance, docking scores (Friesner et al., 2004; Trott & Olson, 2010) lack sufficient accuracy, while gold standard methods like Free Energy Perturbation (FEP) (Zwanzig, 1954) demand enormous computational resources. In recent years, driven by the accumulation of large-scale affinity experimental data and the rapid advancement of deep learning models, e.g. AlphaFold (Jumper et al., 2021), deep learning-based affinity prediction has gradually advanced (Ö ztü rk et al., 2018; Yu et al., 2023; Yang et al., 2024; Wang et al., 2025; Passaro et al., 2025).

Despite these advancements, existing models face a critical challenge in distinguishing *hard inactives* from active ligands. Hard inactives are molecules structurally similar to active compounds but lacking biological activity. We theoretically analyze this limitation in Section 2 and find that it stems from the Lipschitz continuity constraint on neural networks in the Euclidean space. This constraint ensures that model outputs remain close when inputs are sufficiently similar. Consequently, for a known high-affinity molecule, the network predicts high affinity for structurally similar inactives, often ranking them higher than structurally distinct, moderate-affinity molecules. Hard inactives are prevalent in biochemical space, and this *local insensitivity* issue undermines the reliability of rankings and reduces the overall precision of affinity ranking models.

To address this local insensitivity issue, we require a metric space that can break through the Lipschitz continuity constraint. Inspired by the use of hyperbolic space’s negative curvature for modeling hierarchical semantics in multimodal learning (Desai et al., 2023; Pal et al., 2024), we investigate its potential for distinguishing hard inactives. In Section 4.1, we provide a theoretical proof demonstrating that by scoring affinity as the *negative geodesic distance* between targets and small molecules in hyperbolic space, infinitesimal variations on molecule embedding in the target’s tangential direction can break through the Lipschitz limit. Intuitively, the target’s tangent direction should govern global binding factors, such as conformational fit and hydrophobic compatibility, while its radial direction governs local strength factors, such as contact area and bond intensity.

Driven by this geometric intuition, we propose AlphaRank, which projects protein-ligand embeddings into a hyperbolic manifold for joint optimization. The framework employs a weighted ranking loss to differentiate the affinity levels of active molecules, specifically prioritizing pairs that are indistinguishable in Euclidean space. Simultaneously, a geometric constraint separates active molecules from decoys by pushing them into and out of the target’s hyperbolic cone, respectively. Beyond enhancing predictive performance, this joint optimization also yields the additional advantage of a embedding space with strong physical interpretability.

AlphaRank outperforms the state-of-the-art Boltz-2 (Passaro et al., 2025) in both affinity ranking among active molecules and the discrimination of active from inactive compounds. Additionally, quantitative experiments and extensive case studies demonstrate that AlphaRank achieves strong performance on similar molecular pairs that were challenging for previous models to distinguish. Despite being a structure-free approach, our method bridges the gap between traditional structure-based physical and deep learning models, while offering significantly broader applicability.

Our contributions are three-fold:

- We identify the Lipschitz continuity bottleneck in Euclidean space for distinguishing hard inactives and prove that hyperbolic negative geodesic distance effectively breaks this limitation.
- We develop a joint optimization framework that integrates a proximity-aware ranking loss for active molecules with a hyperbolic cone constraint to separate actives from inactives.
- We demonstrate AlphaRank’s state-of-the-art performance in both lead optimization and hit identification, especially in distinguishing between similar molecules.

## 2. The Limitation of Euclidean Metrics

In this section, we analyze affinity ranking from the perspective of function learning and further characterize the mathematical barriers imposed by Euclidean metrics.

Let **x** ∈ ℝ^*m*^ denote the embedding encapsulating protein target information, and **y** ∈ ℝ^*n*^ the embedding encoding ligand information. Our objective is to learn a scoring function *F* : ℝ^*m*^ *×* ℝ^*n*^ → ℝ that preserves the relative order of binding affinities. Formally, for a fixed target **x**, the function must satisfy the ranking consistency property:

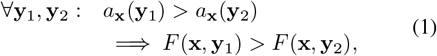

where *a*_**x**_(**y**) denotes the ground-truth binding affinity.

Since **y** is derived from a continuous neural encoder, structural similarities in the input molecules are preserved as Euclidean proximity in the ligand embedding space. Under the Lipschitz continuity constraint, the variation in model predictions is strictly bounded by the distance between these ligand embeddings:

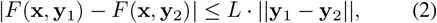

where *L* denotes the Lipschitz constant. This geometric constraint, which compels similar molecular embeddings to yield consistent scores, is a key and reasonable characteristic of neural networks intended to ensure generalization. In the subsequent analysis, we fix the target representation **x**, without loss of generality, treating the scoring function *F* solely as a function of the ligand embedding **y**.

However, such geometric constraints conflict with the biological reality where hard inactives are prevalent. Hard inactives are molecules that are structurally similar to certain active small molecules but lack activity. Due to the Lipschitz constant, neural networks tend to assign scores to hard inactives that are similar to those of their structurally analogous active molecules.

To formally demonstrate the inadequacy of Euclidean metrics in the presence of hard inactives, we formalize a proposition that establishes Lipschitz continuity inevitably results in incorrect ranking outcomes when hard inactives are present.

### Proposition 2.1

(Ranking Failure Induced by Euclidean Lipschitz Continuity). *Let F* ( ·) *be an L-Lipschitz continuous scoring function under the Euclidean metric. For a fixed target, consider three ligands:*

- **y**_1_: *A high-affinity active ligand*.
- **y**_2_: *A moderate-affinity active ligand, where F* (**y**_1_) − *F* (**y**_2_) = Δ *>* 0.
- 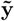: *A hard inactive structurally similar to* **y**_1_, *such that* 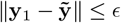. *If the structural difference ϵ is sufficiently small to satisfy L* · *ϵ <* Δ, *the model will inevitably rank the hard inactive higher than the moderate-affinity active ligand* **y**_2_.

This proposition mathematically proves that a stable Euclidean model with *L <* ∞ is intrinsically unable to correctly rank hard inactives against other active ligands, thus necessitating a non-Euclidean geometry for metricizing binding affinity. Therefore, we attempt to find a solution to this problem in Hyperbolic space.

## 3. Preliminaries

To clearly present our method, we first introduce key fundamental concepts of Hyperbolic space in this section.

### Lorentz model and Lorentzian inner product

We adopt the commonly used Lorentz model in hyperbolic space. Under this model, the *n*-dimensional hyperbolic manifold 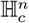 with constant negative curvature *c* (*c >* 0) is defined as the upper sheet of a hyperboloid in the ambient space ℝ^*n*+1^. A point 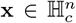 is represented as **x** = [**x**_*space*_, *x*_*time*_]^⊤^, where the spatial component **x**_*space*_ ∈ ℝ^*n*^ and time component *x*_*time*_ ∈ ℝ satisfy the manifold constraint ⟨**x, x** ⟩_ℒ_ = −1*/c* with *x*_*time*_ *>* 0. The geometry is governed by the Lorentzian inner product:

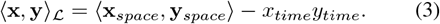

### Geodesic Distance

For any two points 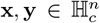, the geodesic distance represents the length of the shortest path between them on the manifold, induced by the inner product:

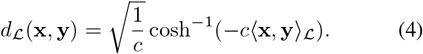

### Exponential Mapping

The exponential mapping bridges Euclidean and hyperbolic spaces by leveraging the tangent space at the origin 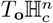 as a local Euclidean approximation. Specifically, a tangent vector **v** (where *v*_*time*_ = 0) is projected onto the manifold as follows:

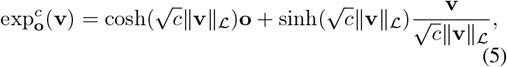

where 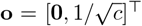 denotes the origin. This operation provides a differentiable mechanism to map Euclidean embeddings from the encoder directly into hyperbolic space.

### Hyperbolic Entailment Cones

While not intrinsic to hyperbolic geometry, hyperbolic entailment cones were introduced to model hierarchical partial orders (Vendrov et al., 2015; Ganea et al., 2018), particularly in multimodal learning where general textual concepts subsume specific visual instances (Pal et al., 2024). For a point **x**, the entailment cone C_**x**_ is defined by a half-aperture angle *ω*(**x**) (Le et al., 2019), which inversely scales with the distance from the origin:

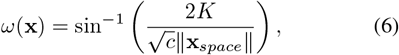

where *K* = 0.1 is a boundary constant. Entailment is then determined by the hyperbolic exterior angle *ϕ*_ℋ_ (**x, y**), representing the deviation *π* − ∠**oxy**:

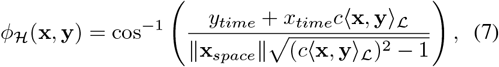

and **y** is entailed by **x** if and only if *ϕ*_*H*_(**x, y**) ≤ *ω*(**x**).

## 4. Method

### 4.1. Breaking Lipschitz Continuity in Hyperbolic Space

To address the ranking failure identified in Proposition 2.1, we propose leveraging the Lorentz model to overcome the limitations imposed by Lipschitz continuity constraints.

In practice, the raw embeddings **x** and **y** in Sec 2 originate from pre-trained encoders, which presents two concerns: their dimensions are often mismatched, and their fixed representations lack the flexibility required for task-specific ranking. To resolve this, we employ learnable projection networks to map these inputs into a unified tangent space, yielding the task-specific feature vectors **p** and **m**. Once lifted onto the manifold, binding affinity is scored as the *negative geodesic distance* between these hyperbolic embeddings. We now analyze how this geometric formulation enables the model to break local Lipschitz constraints.

#### Proposition 4.1

(Breaking Lipschitz Constraints via Hyperbolic Tangential Amplification). *Let p* ∈ ℝ^*n*^ \ {**0**}*and m* ∈ ℝ^*n*^ \ {**0**} *denote the protein and molecule embedding, respectively. We assume that p and m are linearly independent (non-collinear). Without loss of generality, we assume the hyperbolic space has unit negative curvature* −1. *We define the exact hyperbolic affinity scoring function as:*

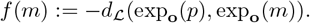

*Consider a perturbation* Δ*s* ∈ ℝ^*n*^, *decomposed relative to the target radial axis p:*

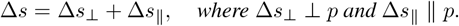

*Provided that* Δ*s*_⊥_ *has a non-vanishing projection onto the tangential subspace of m (i.e*., 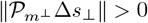*), for sufficiently small* ∥Δ*s*∥, *the affinity variation is strictly bounded below by:*

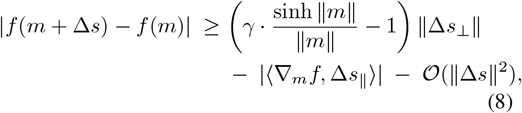

*where γ >* 0 *is a geometric constant defined by the alignment between* Δ*s*_⊥_ *and the tangential component of the geodesic direction:*

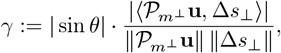

*with* **u** *denoting the unit Riemannian tangent vector of the geodesic from* exp_**o**_(*m*) *to* exp_**o**_(*p*), *and θ the angle between this geodesic and the radial axis m*.

This proposition formalizes that small perturbations along the tangential direction of *m* can induce large changes in the hyperbolic affinity, which leads to the corollary below.

#### Corollary 4.2

(Non-Lipschitz Nature of Hyperbolic Scoring). *The hyperbolic affinity scoring function f* (*m*) *is not Lipschitz continuous with respect to the Euclidean metric on* ℝ^*n*^. *Specifically, there exists no finite constant K* ≥ 0 *such that* |*f* (*m*_1_) − *f* (*m*_2_)| ≤ *K*∥*m*_1_ − *m*_2_∥ *for all m*_1_, *m*_2_ ∈ ℝ^*n*^.

#### Proposition 4.3

(Persistence of Non-Lipschitzness through Neural Networks). *Let f* : ℝ^*n*^ → ℝ *denote the hyperbolic affinity scoring function defined in Proposition 4.1. Let g* : ℝ^*d*^ → ℝ^*n*^ *be a Lipschitz continuous map. Assume that:*

1. ***Unboundedness:*** *The image g*(ℝ^*d*^) *is unbounded;*
2. ***Non-vanishing Projected Jacobian:*** *There exists a sequence* { *y*_*k*_} *such that* ∥ *g*(*y*_*k*_) ∥ → ∞, *and the Jacobian J*_*g*_(*y*_*k*_) *maintains a non-vanishing projection onto the tangential direction of maximal growth of f*. *Specifically, if* 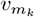 *denotes the unit tangential direction of* ∇*f* (*g*(*y*_*k*_)), *then:*

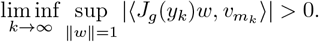

*Then the composition F* (*y*) = *f* (*g*(*y*)) *is not Lipschitz continuous with respect to the Euclidean metric*.

This result theoretically validates that our proposed scoring function *F* effectively breaks the Euclidean Lipschitz constraint. By inducing non-Lipschitz behavior via the hyperbolic manifold, the model gains the necessary geometric capacity to model sharp affinity changes, particularly in challenging *hard inactive* scenarios.

**Biological Interpretation.** Proposition 4.1 demonstrates a tangential dominance in affinity scoring, aligning closely with established biological principles. To better convey this intuition, we set **p** as the origin and primary direction, such that **m**^*′*^ = **m** − **p** represents the coordinates of **m** within this localized reference frame. Here, the local perturbation Δ**s**, corresponds to the differential variation of **m**^′^. This vector **m**^′^ can then be decomposed into orthogonal components 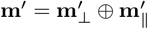, relative to the direction of *p*. The *tangential component* 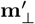 exerts a super-Lipschitz influence on the affinity score, to capture global binding fitness factors such as conformational complementarity and electrostatic matching. Minimizing this tangential component 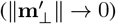 is equivalent to maximizing the binding fitness score, e.g. a judgement of precise conformational match. In contrast, the *radial component* 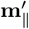 maintains linear stability and Lipschitz continuity to represent local binding strength factors including the contact area and bond intensity. Minimizing this radial residual 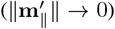 signifies maximizing the binding strength socre.

### 4.2 Model Architecture

We employ a pretrained co-folding model to generate embeddings for both proteins and small molecules. This approach offers two main advantages. First, co-folding models require only the protein sequence and the molecular SMILES as input, eliminating the need for precise 3D binding or docked structures. This significantly broadens the applicability of our method. Second, the capability of co-folding models, as demonstrated by frameworks like AlphaFold3, in modeling molecular interactions is well-established, allowing us to leverage proven interaction modeling for affinity ranking without training from scratch. Specifically, we use Protenix (Team et al., 2025) as our feature encoder. The protein embedding **x** is obtained by inputting the target protein sequence alone. The molecule embedding **y** is derived by inputting both the protein sequence and the molecular SMILES, as we posit that a molecule’s properties are context-dependent and its features are more informative when conditioned on the specific target protein. Details of this process are provided in Appendix D.

To project these features into hyperbolic space, we first align them to a tangent space via two parameterized projectors: **p** = *g*_*P*_ (**x**) and **m** = *g*_*L*_(**y**). In practice, our parameterized projectors theoretically satisfies the conditions in Proposition 4.3,see implementation details in appendix E. An exponential mapping then projects them into the hyperbolic space: 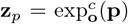 and 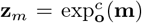. Finally, the binding affinity score is scored as the negative geodesic distance between **z**_*p*_ and **z**_*m*_.

### 4.3. Training Objectives

To structure the hyperbolic space described in Section 4.1, which captures global binding fitness factors tangentially and local binding strength factors radially, we employ two training objectives: a ranking loss to order active molecules by their geodesic distance to the protein, and a geometric loss to enforce cone separation between actives and inactives. Training data is derived from ChEMBL35 (Zdrazil et al., 2024) and structured as lists of the form 𝒯 = {*t*_1_, *t*_2_, …, *t*_*k*_, *t*_*decoy*_}. Each list contains *k* active ligands sorted by decreasing affinity, with significant affinity gaps to ensure deterministic supervision. Since experimentally validated inactive data is often scarce in public databases, we augment our supervision with a decoy molecule *t*_*decoy*_, obtained via cluster-based negative sampling. See Appendix C for detailed data preparation protocols. Specifically, for each ligand *t*_*i*_ ∈ 𝒯, let **y**_*i*_ denote the molecular embedding derived from the Protenix encoder. This yields **m**_*i*_ = *g*_*L*_(**y**_*i*_), and its hyperbolic projection 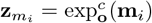.

#### 4.3.1. Geodesic-based Proximity-aware Ranking

Our choice of a ranking loss over regression is motivated by its robustness, since relative ordering in data is more consistent across assays than absolute affinity values, which are prone to experimental noise. A primary challenge of ranking is distinguishing molecules with similar embeddings in Euclidean space but divergent binding affinities. To address this, we introduce a weighting coefficient *ψ*_*ij*_, that scales inversely with the Euclidean distance between embeddings:

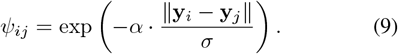

This coefficient forces the separation of proximal pairs on the hyperbolic manifold. The hyperparameter *σ >* 0 provides a normalization scale, while *α >* 0 dictates the focus intensity.

We model the conditional probability of the preference relation *t*_*i*_ *≻ t*_*j*_ using a modulated Sigmoid function *σ*( ·). By incorporating the score margin Δ_*i,j*_ = *F* (**y**_*i*_) − *F* (**y**_*j*_),where *F* (·) is the affinity scoring function defined in Section 4.1. The pairwise loss simplifies to a scaled Softplus form:

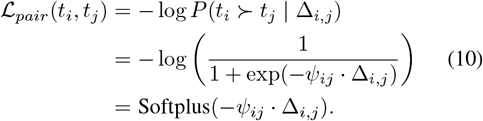

The complete rank loss aggregates these pairwise terms:

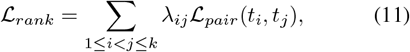

where *λ*_*ij*_ represents the hyperparameter controlling the weight of the pair (*t*_*i*_, *t*_*j*_). In our practical implementation, we set *k* = 3, instantiating the training instance as an ordered triplet. Crucially, this triplet-based optimization is not merely a local heuristic but provides rigorous global guarantees. As formally proven in Appendix C, minimizing this objective is mathematically equivalent to optimizing a global risk over the entire list structure, ensuring that the learned hyperbolic geometry effectively enforces the precise ranking order of high-affinity leads.

#### 4.3.2. Geometric Cone-based Separation

Clearly separating active and inactive molecules within a geometric cone requires more than simple inclusion or exclusion. We therefore introduce an explicit tangential force to drive their representations apart. While implementing such a force directly in hyperbolic space is complex, we achieve an equivalent effect by optimizing the *Euclidean alignment angle θ*, defined as the angle between protein and molecule embeddings in the tangent space. Proposition 4.4 validates this approach by establishing *θ* as a proxy for the hyperbolic exterior angle *ϕ*_*H*_.

##### Proposition 4.4

(Monotonic Relation of Euclidean and Hyperbolic Angles). *For a fixed target norm R* = ∥**p**∥, *under the entailment assumption (*∥**m**∥ *>* ∥**p**∥*), the hyperbolic exterior angle ϕ*_*H*_ *is a strictly monotonically increasing function of the Euclidean alignment angle θ*.

This monotonicity implies that to achieve geometric separation, we should minimize *θ* for active molecules to keep them within the cone, while maximizing it for inactives to push them outside. For a training list 𝒯, we employ a RankNet-style loss to maximize the cosine similarity of the active ligands against a decoy:

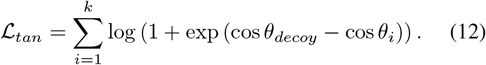

Here, *θ*_*decoy*_ and *θ*_*i*_ denote the Euclidean alignment angles between the target vector **p** and the feature vectors of the decoy **m**_*decoy*_ and the *i*-th active ligand **m**_*i*_, respectively.

As derived in Appendix B, this tangent space separation in the Euclidean space guarantees a ranking margin that scales linearly with the target norm *R* in the large-radius regime, effectively translating subtle angular differences into significant geodesic distances.

While the separation loss described above does not guarantee that active ligands are placed inside the target cone, we additionally minimize an *entailment loss* (Le et al., 2019; Desai et al., 2023) to ensure that all active ligands are compressed into the specific hyperbolic cone 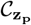:

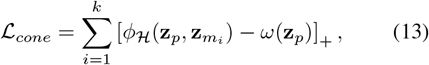

where [·]_+_ = max(0, ·) denotes the ReLU operator.

The complete geometric cone based separation loss combines both constraints:

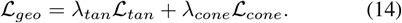

**Biological Interpretation of Cone Separation.** We find that translating the cone separation constraint back into the Euclidean tangent space, the imposed constraint aligns with biological intuition. By analyzing the **p**-localized ligand representation **m**^*′*^ = **m** − **p**, we identify a Euclidean funnel-like region, which we term the *tolerance funnel*, defined by the coefficient *ω*(*R*). This coefficient corresponds to the half-aperture angle in Equation 6. Since it depends only on the norm *R* = ∥**p**∥ of the protein embedding, we denote it here as *ω*(*R*).

##### Proposition 4.5

(The Euclidean Tolerance Funnel Constraint). *Let* **m**^*′*^ *be decomposed into tangential and radial components* 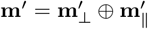. *Driven by the cone separation, if an active ligand is successfully entailed within the cone, i.e*. 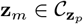, *then its tangent components satisfy the strict linear convergence constraint:*

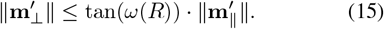

This inequality is consistent with the established physical principles of molecular recognition (Chothia & Janin, 1975; Mobley & Dill, 2009). In the high-affinity regime, the tight packing of the ligand causes 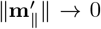, necessitating a simultaneous reduction in 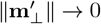 and leaving no margin for global binding fitness error. Essentially, exceptional binding strength becomes inseparable from near-perfect fitness. In contrast, the constraint relaxes proportionally for low-strength interactions where larger values of 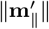 permit greater conformational flexibility without incurring severe physical penalties.

#### 4.3.3. Final Loss

The complete training objective is a weighted sum of the ranking loss and the geometric loss:

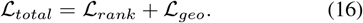

## 5. Experiments

### 5.1. Experiment Setup

#### Datasets

To evaluate AlphaRank’s ability to rank molecular affinities, we benchmark it on the widely used lead optimization datasets, JACS (Wang et al., 2015) and Merck (Schindler et al., 2020). Its capability to distinguish actives from inactives is further assessed on the standard hit identification dataset CYP3A4 (Veith et al., 2009). The model is trained on data constructed from ChEMBL35, and is deduplicated using canonical sequences from UniProt to prevent data leakage.

#### Baselines

We compare AlphaRank against a comprehensive suite of existing approaches spanning four distinct modeling paradigms. The evaluation encompasses *Physics-based methods*, including the rigorous Free Energy Perturbation (Wang et al., 2015) and MM-GB/SA (Genheden & Ryde, 2015), alongside *Sequence-based models* such as DeepDTA (Ö ztürk et al., 2018), DeepPurpose (Huang et al., 2020), and BIND (Lam et al., 2024) that rely solely on protein sequences and ligand SMILES. We also incorporate *Structure-based models*, incuding PBCNet (Yu et al., 2023), EHIGN (Yang et al., 2024), GET (Kong et al., 2024),

BindNet (Feng et al., 2024), and LigUnity (Feng et al., 2025), which requires pocket structures, and *Co-folding-based models* represented by Boltz-2 (Passaro et al., 2025) and Protenix-ipTM (Team et al., 2025). To ensure fairness, all baselines are evaluated using their official implementations or following reported protocols. Details of the AlphaRank model implementation and training are provided in Appendix E.

### 5.2. Results on Lead Optimization

Table 1 demonstrates that AlphaRank achieves superior performance on both the JACS and Merck datasets. It consistently outperforms other methods that are structure-free, including sequence-based and co-folding-based methods. Furthermore, AlphaRank surpasses existing structure-based deep learning models and approaching to the gold-standard FEP+. These results confirm that mapping embedding into hyperbolic space and scoring affinity with negative geodesic distance achieve efficient affinity ranking.

**Figure 1.**
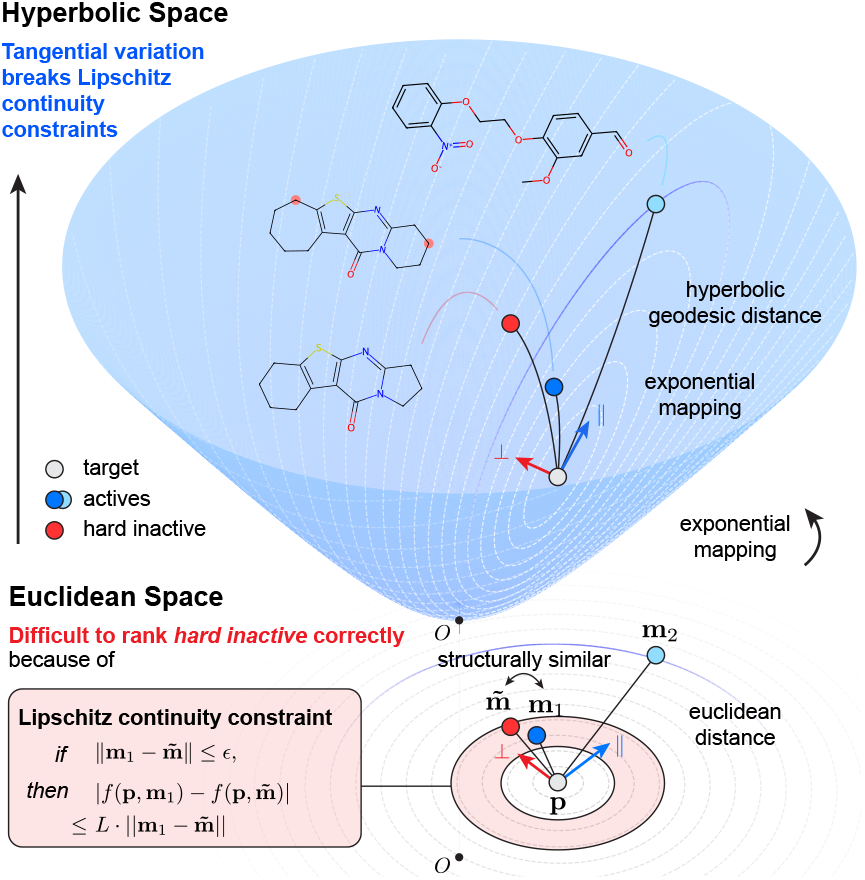
Comparison of affinity ranking in Euclidean vs. Hyperbolic spaces. While Euclidean models often struggle to distinguish *hard inactives* from actives due to Lipschitz-induced insensitivity (bottom), AlphaRank measures affinity with hyperbolic geodesic distance, enabling subtle tangential variations to break Lipschitz continuity constraints for precise ranking (top).

**Figure 2.**
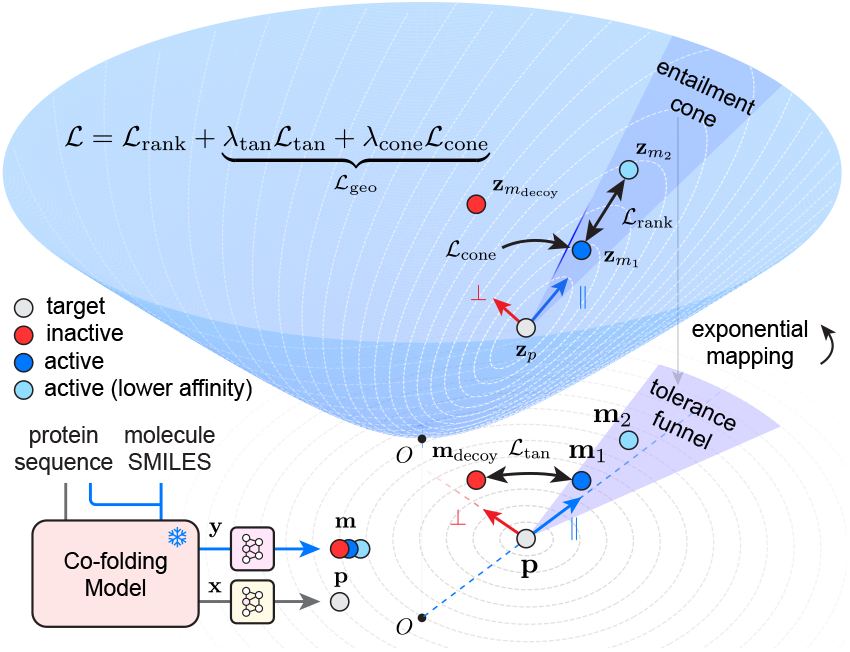
AlphaRank’s architecture and training objectives.

**Table 1.** Lead Optimization performance comparison on the JACS and Merck benchmarks. Metrics *r* and *ρ* denote the Pearson and Spearman correlation coefficients, respectively.

| Type | Model | JACS |  | Merck |  |
| --- | --- | --- | --- | --- | --- |
| | | $r$ | $\rho$ | $r$ | $\rho$ |
| Physics | FEP+ | 0.7811 | 0.7595 | 0.6960 | 0.6798 |
|  | MM-GB/SA | 0.1489 | 0.2011 | 0.1299 | 0.1299 |
| Sequence | DeepDTA | 0.3485 | 0.4098 | 0.3243 | 0.2978 |
|  | DeepPurpose | -0.0306 | -0.0313 | 0.0375 | 0.0381 |
|  | BIND | 0.4816 | 0.4750 | 0.3289 | 0.2947 |
| Structure | PBCNet | 0.3939 | 0.3799 | 0.4058 | 0.4075 |
|  | EHIGN | 0.5787 | 0.5814 | 0.4246 | 0.3830 |
|  | GET | 0.4034 | 0.3753 | 0.4203 | 0.4214 |
|  | BindNet | 0.5481 | 0.5368 | 0.4037 | 0.3477 |
|  | LigUnity | 0.5705 | 0.5774 | **0.5323** | 0.4994 |
| Co-folding | Boltz-2 | 0.5231 | 0.5285 | 0.4298 | 0.4013 |
|  | AlphaRank | **0.6461** | **0.6536** | 0.5243 | **0.5327** |

**Table 2.** Performance comparison across edges in JACS and Merck benchmarks.

| Type | Model | JACS |  | Merck |  |
| --- | --- | --- | --- | --- | --- |
|  |  | Acc | AUC | Acc | AUC |
| Physics | FEP+ | 0.7806 | 0.8515 | 0.7496 | 0.8545 |
|  | MM-GB/SA | 0.6396 | 0.7182 | 0.5212 | 0.5607 |
| Sequence based | DeepDTA | 0.5996 | 0.6378 | 0.5632 | 0.5812 |
|  | DeepPurpose | 0.5039 | 0.4489 | 0.4922 | 0.4962 |
|  | BIND | 0.3828 | 0.3254 | 0.4660 | 0.4318 |
| Structure based | PBCNet | 0.6371 | 0.6886 | 0.6083 | 0.6239 |
|  | EHIGN | 0.6436 | 0.7329 | 0.6092 | 0.6563 |
|  | LigUnity | 0.6694 | 0.7737 | 0.6176 | 0.7009 |
| Co-folding based | ipTM | 0.5679 | 0.6406 | 0.5547 | 0.6058 |
|  | Boltz-2 | 0.6793 | 0.7863 | 0.6796 | 0.7431 |
|  | AlphaRank | **0.7038** | **0.8089** | **0.7059** | **0.7627** |

Beyond the overall results, we further examined the performance of each method on hard pairs, which are referred to as *edges* in the JACS and Merck, where each edge corresponds to a chemically meaningful ligand pair selected based on structural similarity, synthetic feasibility and SAR significance. These pairs reflect realistic decisions in lead optimization pipelines where precise molecular ranking is more impactful. Results in Table 1 show that our method outperforms all existing approaches by a large margin and approaches FEP+. Its excellent performance on these structurally similar small molecule pairs proves that modeling in hyperbolic space effectively helps the model distinguish affinity changes caused by minor ligand differences.

### 5.3. Results on Hit Identification

CYP3A4 dataset is specifically designed for evaluating hit identification and includes activity cliff data of Cytochrome P450 3A4 inhibitors and substrates experiment results. Results in Table 3 demonstrate that AlphaRank outperforms other models in hit identification. It not only distinguishes small molecules with varying affinities but also exhibits strong capability in differentiating actives from inactives. In contrast, Boltz-2 employs two separate predictors. One is a binary classifier for active-inactive distinction and the other a regression model for absolute affinity prediction. These two predictors show significant inconsistency on this task with the regression head performing even better. This further validates the challenge of distinguishing inactive in Euclidean space, especially when the inactive is similar to the compared active, leading to model inconsistency. By contrast, AlphaRank learns a single, coherent embedding space by jointly optimizing both geodesic distance ranking and hyperbolic cone separation. As previously explained in Section 4.1, this enables the model to capture global binding fitness primarily in the tangential direction and binding strength in the radial direction within the same unified representation, resulting in a highly consistent space.

**Table 3.** Performance comparison on the CYP3A4 benchmark.

| Type | Model | AUC |
| --- | --- | --- |
| Sequence | DeepDTA | 0.6650 |
|  | DeepPurpose | 0.4906 |
|  | BIND | 0.7205 |
| Co-folding | Boltz-2 (binary) | 0.6112 |
|  | Boltz-2 (affinity) | 0.7263 |
|  | AlphaRank | **0.7589** |

### 5.4. Ablation Studies

Table 4 shows the results of ablation studies to evaluate the contribution of each core component within AlphaRank. Removing any individual module leads to a marked drop in performance, with Pearson *r* falling from 0.524 to between 0.41 and 0.46. Specifically, replacing the hyperbolic manifold with a Euclidean counterpart (w/o hyp) results in the most severe drop, confirming the necessity of employing non-Euclidean geodesic metrics to accurately model binding affinity. The significant drops upon removing ℒ_*tan*_ or ℒ_*cone*_ indicate that separating actives from decoys in the tangent space and confining actives within the hyperbolic cone are complementary for refining the embedding space. Finally, disabling the proximity-aware weighting (w/o *ψ*_*ij*_) highlights the importance of amplifying gradients for ligand pairs that are spatially close in the tangent space, thereby enforcing their separation on the hyperbolic manifold.

**Table 4.** Ablation results on the Merck benchmark.

| Setting | Module |  |  | Merck |  |
| --- | --- | --- | --- | --- | --- |
| | $\psi_{ij}$ | $\mathcal{L}_{tan}$ | $\mathcal{L}_{cone}$ | $r$ | $\rho$ |
| Full model | ✓ | ✓ | ✓ | 0.5243 | 0.5327 |
| w/o hyp | ✗ | ✓ | ✗ | 0.4189 | 0.4093 |
| w/o $\mathcal{L}_{tan}$ | ✓ | ✗ | ✓ | 0.4699 | 0.4485 |
| w/o $\mathcal{L}_{cone}$ | ✓ | ✓ | ✗ | 0.4434 | 0.4313 |
| w/o $\psi_{ij}$ | ✗ | ✓ | ✓ | 0.4564 | 0.4538 |

### 5.5. Case study

Through concrete case studies, we find that AlphaRank significantly outperforms existing models on challenging inactive cases. Table 5 details a representative example from PubChem AID 588549, comparing Boltz2 and AlphaRank, while further examples are provided in Appendix F. Despite sharing high structural similarity with an ECFP4 Tanimoto similarity of 0.9, these two molecules possess opposite labels. Boltz-2 fails to discriminate between the active and hard inactive molecules. Notably, Boltz-2 assigns a higher affinity score to the inactive compound and predicts a binding pose that suggests even stronger protein-ligand interactions than those of the active counterpart. In contrast, AlphaRank captures the critical structural nuances that differentiate them. Embedding analysis reveals that the hard inactive molecule introduces a significant orthogonal deviation 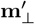 relative to the protein target, with the norm of 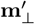 increasing fourfold from 0.38 to 1.54. By leveraging non-Euclidean geodesic metrics, AlphaRank translates this geometric shift into a scoring margin of Δ = +4.48. This demonstrates a significant advantage in reducing false positives arising from hard inactives.

**Table 5.**
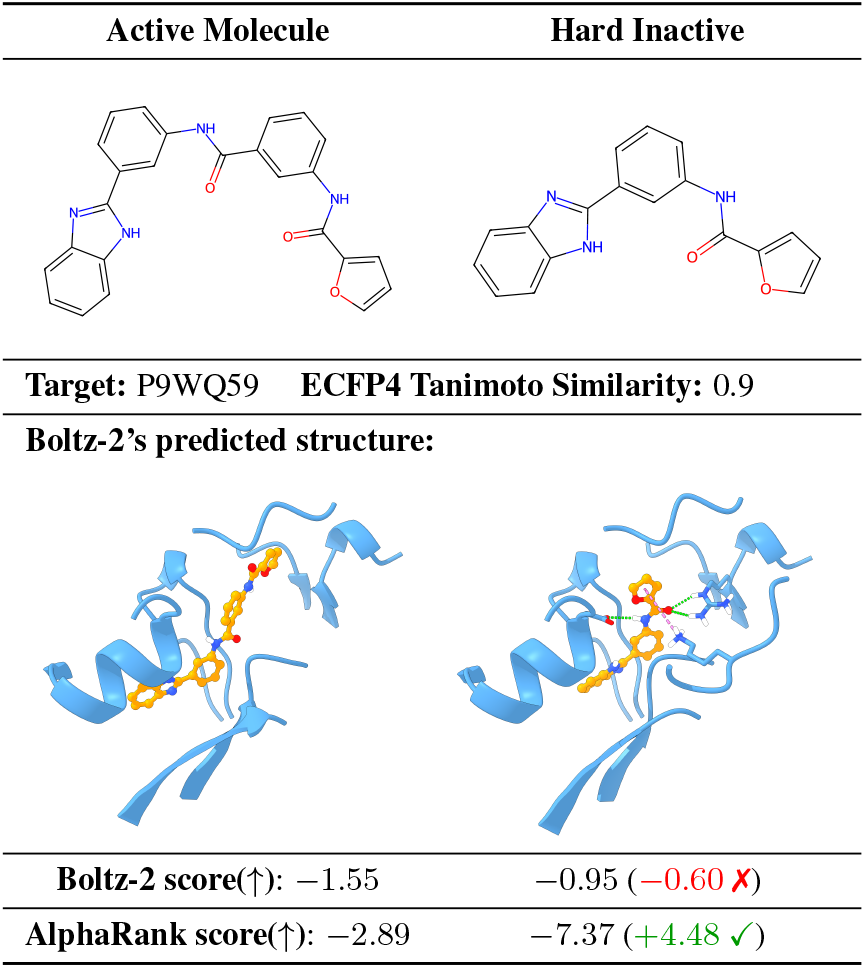
AlphaRank and Boltz-2’s performance on a typical *hard inactive* case.

| Active Molecule | Hard Inactive |
| --- | --- |
| Target: P9WQ59 ECFP4 Tanimoto Similarity: 0.9 |  |
| Boltz-2’s predicted structure: |  |
| Boltz-2 score(↑): -1.55 | -0.95 (-0.60 ×) |
| AlphaRank score(↑): -2.89 | -7.37 (+4.48 ✓) |

## 6. Related Works

Traditional physics-based methods for protein–ligand binding affinity prediction vary in fidelity. Molecular docking evaluates geometric and physicochemical compatibility of mostly static complexes, while molecular dynamics integrates conformational flexibility and temporal evolution. Free energy perturbation such as FEP+ (Wang et al., 2015) achieves near-gold-standard accuracy by explicitly accounting for atomic interactions, but its prohibitive computational cost restricts application to small libraries and late-stage optimization. Early deep learning models relied on sequence-based 1D representations of proteins and ligands (Ö ztü rk et al., 2018; Huang et al., 2020; Lam et al., 2024), lacking explicit structural context. Subsequent structure-based approaches incorporated 3D information via graph neural networks and equivariant Transformers on experimentally determined complexes (Yu et al., 2023; Yang et al., 2024; Feng et al., 2024; Kong et al., 2024; Feng et al., 2025), improving accuracy but limiting use to targets with known high-resolution structures. Recent co-folding-based foundation models like Boltz-2 (Passaro et al., 2025) bridge this gap by jointly inferring protein–ligand geometry and affinity from sequence inputs, and our work follows this paradigm to maximize modelable targets while retaining such structure-aware capabilities.

Hyperbolic space has emerged as a powerful manifold for representing data with inherent hierarchical or tree-like organization, owing to its exponential volume growth that preserves hierarchy with low distortion (Nickel & Kiela, 2017; Chamberlain et al., 2017). Foundational works demonstrated that Poincaré or Lorentz embeddings significantly outperform Euclidean approaches in preserving the structure of taxonomies and complex graphs (Ganea et al., 2018). More recent studies have explored multimodal training in hyperbolic space, enabling vision–language models to better capture hierarchical semantics (Desai et al., 2023; Pal et al., 2024). In the context of binding affinity, recent work includes HypSeek (Wang et al., 2025), a structure-based model that utilizes explicit pocket structures in hyperbolic space for virtual screening and affinity ranking.

## 7. Conclusion

This paper highlights the critical yet overlooked hard inactive problem in affinity ranking. We demonstrate that Lipschitz continuity constraints in Euclidean space fundamentally restrict a model’s ability to distinguish hard inactives, and that scoring binding affinity as the negative geodesic distance in hyperbolic space can break this limitation. Based on this theory, we introduce AlphaRank, a framework jointly optimize geodesic proximity aware ranking and geometric cone-based separation objectives. The theoretical foundation of AlphaRank ensures that the learned embedding space is highly interpretable and aligns closely with biological intuition, such as the tangential-radial decomposition of binding factors and the tolerance funnel discussed, thereby enhancing the reliability of the model’s predictions. AlphaRank achieves state-of-the-art performance on widely used lead optimization and hit identification benchmarks, approaching the accuracy of the gold-standard FEP+. Through extensive ablation studies and dedicated case studies on hard inactives, we validate the effectiveness and superiority of modeling affinity ranking in hyperbolic space.

## Impact Statement

This paper presents work whose goal is to advance the field of machine learning-based affinity ranking. There are many potential societal consequences of our work, none of which we feel must be specifically highlighted here.

## A. Proofs of Propositions

*In this section, we provide rigorous proofs for the propositions and corollaries presented in the main text*.

### Proof of Proposition A.1

Let F ( ·) be an L-Lipschitz continuous scoring function under the Euclidean metric. Consider three ligands for a fixed target:

- **y**_1_: *A high-affinity active ligand*.
- **y**_2_: *A moderate-affinity active ligand, where F* (**y**_1_) − *F* (**y**_2_) = Δ *>* 0.
- 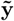: *A hard inactive structurally proximate to* **y**_1_, *such that* 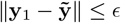.

If the structural difference ϵ is sufficiently small such that *L* · ϵ < Δ, then the model must incorrectly rank the inactive higher than the moderate active ligand ***y***_*2*_.

*Proof*. By the definition of Lipschitz continuity, the score difference between the high-affinity active **y**_1_ and inactive 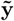 is bounded by:

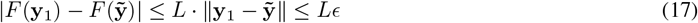

Expanding the absolute value inequality, we have a lower bound for the inactive’s score:

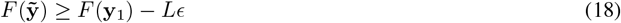

Since we assume the active gap is large, i.e., *F* (**y**_1_) = *F* (**y**_2_) + Δ, we substitute this into the inequality:

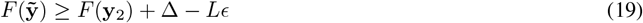

Given the condition Lϵ < Δ (which holds for hard inactives where ϵ → 0 and bounded *L*), it strictly follows that Δ − *Lϵ* > 0. Therefore:

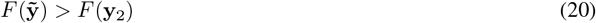

This concludes the proof. The model assigns a higher affinity score to the inactive 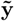 than to the true active **y**_2_, violating the Ranking Consistency.

### Proof of Proposition 4.1

#### Proposition A.2

*We identify the Euclidean feature space* ℝ^*n*^ *with the tangent space at the origin T*_***o***_ℍ^*n*^ *via the logarithmic map.Let p* ∈ ℝ^*n*^ *\* {***0***} *and m* ∈ ℝ^*n*^ *\* {***0***} *denote the feature representations encapsulating the protein and ligand information, respectively. We assume that p and m are linearly independent (non-collinear). Without loss of generality, we assume the hyperbolic space has unit negative curvature* −*1. We define the exact hyperbolic affinity scoring function as*:

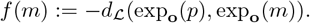

Consider a perturbation Δs ∈ ℝ^*n*^, decomposed relative to the target radial axis *p*:

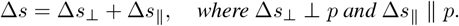

*Provided that* Δ*s*_⊥_ *has a non-vanishing projection onto the tangential subspace of m (i.e*., 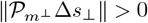*), for sufficiently small* ∥Δ*s*∥, *the affinity variation is strictly bounded below by:*

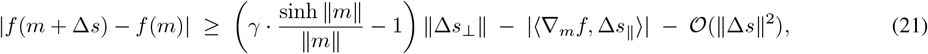

where γ > *0* is a geometric constant defined by the alignment between Δs_⊥_ and the tangential component of the geodesic direction:

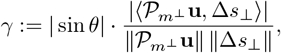

with **u** denoting the unit Riemannian tangent vector of the geodesic from *exp*_***o***_(m) to *exp*_***o***_(p), and θ the angle between this geodesic and the radial axis m.

***Proof***. Gradient Structure and Anisotropic Scaling.

Let Φ(*x*) = exp_**o**_(*x*). By the chain rule, the Euclidean gradient of the affinity function is:

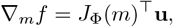

where *J*_Φ_(*m*) is the Jacobian of the exponential map at *m*. It is a standard result in hyperbolic geometry that the Jacobian of the exponential map has eigenvalues 1 (along the radial direction) and 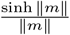 (along tangential directions). Thus, *J*_Φ_(*m*) admits an anisotropic spectral decomposition: it preserves radial vectors but expands tangential vectors by the conformal factor 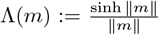. Let 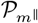 and 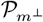 be the orthogonal projectors onto the radial and tangential subspaces of *m*.

The Euclidean gradient decomposes as:

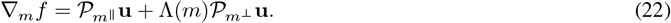

Due to the linear independence assumption, 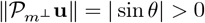.

## Linearization and Decomposition

For sufficiently small ∥Δ*s*∥, the Taylor expansion gives:

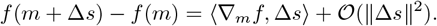

Substituting Δ*s* = Δ*s*_⊥_ + Δ*s*_∥_, we have:

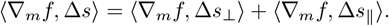

## Rigorous Lower Bound via Reverse Triangle Inequality

Applying the reverse triangle inequality |*A* + *B*| ≥ |*A*| − |*B*|:

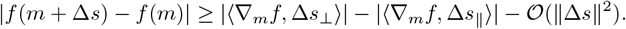

We analyze the signal term |⟨∇_*m*_*f*, Δ*s*_⊥_⟩| rigorously. Substituting Eq. (22):

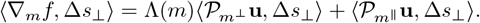

Applying the reverse triangle inequality again to separate the amplified tangential component from the non-amplified radial projection:

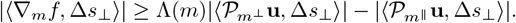

Using 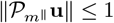 (since **u** = 1), the term 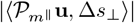 is strictly bounded by ∥Δ*s*_⊥_ ∥. Meanwhile, substituting the definition of *γ*:

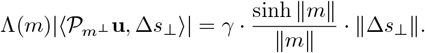

Thus, we obtain the strict lower bound for the signal term:

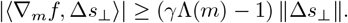

## Conclusion

Substituting this back into the main inequality yields:

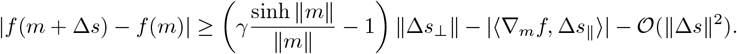

This completes the proof.

*Remark* A.3. The lower bound in Proposition 4.1 is asymptotic in nature. For bounded ∥*m*∥, the coefficient (*γ*Λ(*m*) − 1) may be non-positive; however, in the deep embedding regime relevant to our setting, Λ(*m*) grows exponentially and the tangential term strictly dominates all linear contributions.

## Proof of Corollary 4.2

### Corollary A.4

*The hyperbolic affinity scoring function f* (*m*) *is not Lipschitz continuous with respect to the Euclidean metric on* ℝ^*n*^ *(for n* ≥ 2*). Specifically, there exists no finite constant K* ≥ 0 *such that:*

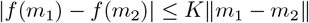

*for all m*_1_, *m*_2_ ∈ ℝ^*n*^.

*Proof*. We proceed by contradiction. Assume that *f* (*m*) is Lipschitz continuous with a Lipschitz constant *K* ≥ 0. By Rademacher’s theorem, a Lipschitz function is differentiable almost everywhere, and its gradient norm is bounded by *K* at differentiable points. Specifically, for any *m* where *f* is differentiable and any unit vector *v* ∈ ℝ^*n*^, the directional derivative must satisfy:

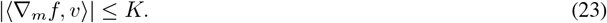

We construct a sequence of inputs that violates this bound. Since the set of non-differentiable points has Lebesgue measure zero, we explicitly construct our sequence {*m*_*k*_} within the domain of differentiability of *f*.

Consider a fixed target *p*. We construct a sequence 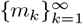 along a fixed ray emanating from the origin that is distinct from the ray passing through *p*. Specifically, let *m*_*k*_ = *k* · *v*_*dir*_ for some unit vector *v*_*dir*_ ∦ *p*. In this setup, as ∥*m*_*k*_∥ → ∞, the angle *θ*_*k*_ between the radial axis *m*_*k*_ and the geodesic to *p* converges to the Euclidean angle between the two rays, which is strictly non-zero by construction. Consequently, the geometric factor is strictly bounded away from zero: *γ*_*k*_ ≥ *c >* 0 for all sufficiently large *k*.

For each *m*_*k*_, we choose a specific perturbation direction Δ*s* to exploit the tangential dominance established in Proposition 4.1:

- We select Δ*s* to be strictly orthogonal to the target radial axis *p* (i.e., Δ*s* ⊥ *p*, implying Δ*s*_∥_ = 0).
- We ensure Δ*s* has a non-vanishing projection onto the tangential component of the geodesic tangent **u** (i.e., 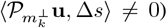. Since *v*_*dir*_ ∦ *p* implies 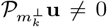, and the ambient dimension satisfies *n* ≥ 2, the orthogonal complement of *p* intersects strictly with the support of the tangential component, guaranteeing that such a direction Δ*s* always exists. (Note: for *n* = 1, the tangential subspace is trivial relative to the radial direction, but high-dimensional embeddings inherently satisfy *n* ≥ 2).

Under this choice, the linear noise term vanishes (∥Δ*s*_∥_∥ = 0).

From Proposition 4.1, specifically Eq. (21), for a sufficiently small ∥Δ*s*∥ *>* 0, the affinity variation is bounded by:

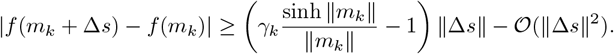

Dividing both sides by ∥Δ*s*∥ and taking the limit ∥Δ*s*∥ → 0, the higher-order term 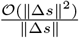 vanishes. We consider the directional derivative along the unit vector 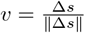. Since *f* is differentiable at *m*_*k*_, this limit equates to the inner product magnitude:

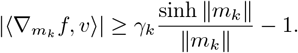

Let 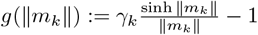. Since sinh(*x*) grows exponentially and *γ*_*k*_ ≥ *c >* 0, we have:

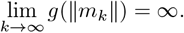

Consequently, for any fixed *K*, there exists an index *k*_0_ such that for all *k > k*_0_, the directional derivative exceeds the bound:

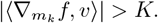

This contradicts the necessary condition for Lipschitz continuity (Eq. (23)). Thus, *f* (*m*) is not Lipschitz continuous with respect to the Euclidean metric.

## Proof of Proposition 4.3

### Proposition A.5

*Let f* : ℝ^*n*^ → ℝ *denote the hyperbolic affinity scoring function defined in Proposition 4.1. Let g* : ℝ^*d*^ → ℝ^*n*^ *be a Lipschitz continuous map. Assume that:*

1. ***Unboundedness:*** *The image g*(ℝ^*d*^) *is unbounded;*
2. ***Non-vanishing Projected Jacobian:*** *There exists a sequence* { *y*_*k*_} *such that* ∥*g*(*y*_*k*_) ∥ → ∞, *and the Jacobian J*_*g*_(*y*_*k*_) *maintains a non-vanishing projection onto the tangential direction of maximal growth of f*. *Specifically, if* 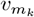 *denotes the unit tangential direction of* ∇*f* (*g*(*y*_*k*_)), *then:*

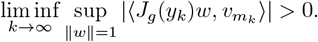

*Proof*. We argue by contradiction.

### Tangential gradient blow-up of *f*

By Proposition 4.1, the gradient of *f* exhibits exponential growth along specific tangential directions as ∥*m*∥ → ∞. Let *m*_*k*_ = *g*(*y*_*k*_) from Assumption 2. There exists a unit tangential direction 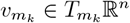 such that

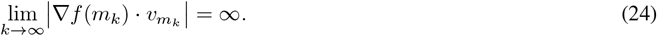

### Compatibility with the lifting map *g*

By Assumption 2, the encoder *g* does not collapse the gradient in the direction of *v*_*m*_. There exists a constant *ε >* 0 such that for all sufficiently large *k*, we can find a direction *w*_*k*_ ∈ ℝ^*d*^ with ∥*w*_*k*_∥ = 1 satisfying

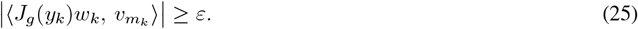

### Tangential–radial decomposition

By the chain rule, ∇*F* (*y*_*k*_) = *J*_*g*_(*y*_*k*_)^⊤^ ∇ *f* (*m*_*k*_). We decompose the gradient of *f* at *m*_*k*_ as

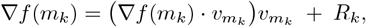

where *R*_*k*_ is the residual component orthogonal to 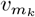. By Proposition 4.1, the tangential component diverges, whereas the residual component *R*_*k*_ remains uniformly bounded. Specifically, there exists a constant *C <* ∞ such that ∥ *R*_*k*_ ∥ ≤ *C* for all *k*.

### Dominance of the tangential contribution

Consider the directional derivative along *w*_*k*_:

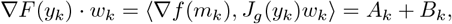

where 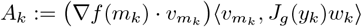 and *B*_*k*_ := ⟨*R*_*k*_, *J*_*g*_(*y*_*k*_)*w*_*k*_⟩.

Applying the reverse triangle inequality:

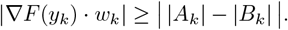

Analyzing the terms as *k* → ∞:

- **Term** *A*_*k*_: By Step 1, 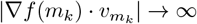, and by Step 2, 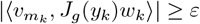. Therefore, |*A*_*k*_| → ∞.
- **Term** *B*_*k*_: Since *g* is Lipschitz, ∥*J*_*g*_(*y*_*k*_)∥ ≤ Lip(*g*) almost everywhere. Combined with the boundedness ∥*R*_*k*_∥ ≤ *C*, we have |*B*_*k*_| ≤ *C* · Lip(*g*) *<* ∞.

Hence, for sufficiently large *k*, the diverging term |*A*_*k*_| dominates the bounded term |*B*_*k*_|, implying

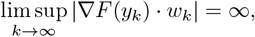

which in turn implies ∥∇*F* (*y*_*k*_)∥ → ∞.

### Violation of Lipschitz continuity

If *F* were Lipschitz continuous with constant *L*, then ∥∇*F* (*y*) ∥ ≤ *L* almost everywhere. This contradicts the unboundedness established in Step 4. Therefore, *F* is not Lipschitz continuous.

## Proof of Proposition 4.4

### Proposition A.6

*For a fixed target norm R* = ∥**p**∥, *under the entailment assumption (*∥**m**∥ *>* ∥**p**∥*), the hyperbolic exterior angle ϕ*_*H*_ *is a strictly monotonically increasing function of the euclidean alignment angle θ*.

*Proof*. Without loss of generality, we assume unit curvature *c* = 1. Let **z**_*p*_ = [**z**_*p,s*_, *z*_*p,t*_]^⊤^ and **z**_*m*_ = [**z**_*m,s*_, *z*_*m,t*_]^⊤^ denote the hyperbolic coordinates of the target and the ligand, respectively. Here, **z**_*p,s*_ and *z*_*p,t*_ represent the spatial and time components of **z**_*p*_. These are derived from the Euclidean tangent vectors **p, m** via the exponential map:

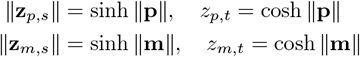

Let *θ* be the Euclidean angle between the tangent vectors **p** and **m**. The Lorentz inner product ⟨**z**_*p*_, **z**_*m*_⟩_*L*_ is given by:

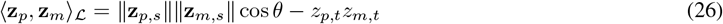

Let *X* = −⟨**z**_*p*_, **z**_*m*_⟩ _ℒ_. Since **z**_*p*_ ≠ **z**_*m*_, we have *X >* 1. The hyperbolic exterior angle *ϕ*_*H*_(**z**_*p*_, **z**_*m*_) at the vertex **z**_*p*_ is defined as:

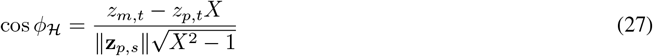

### Derivative of cos *ϕ*_*H*_ **with respect to** cos *θ*

First, we compute the derivative of *X* with respect to cos *θ*:

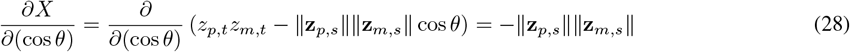

Next, we differentiate cos *ϕ*_*H*_ with respect to *X*:

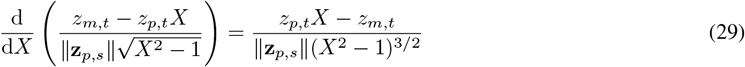

Applying the chain rule:

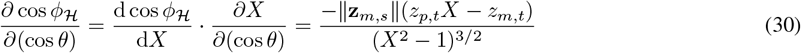

Given the entailment assumption ∥**m**∥ *>* ∥**p**∥, implies that the ligand is further from the origin than the target, ensuring *z*_*m,t*_ *> z*_*p,t*_. Combined with *X >* 1, the term (*z*_*p,t*_*X* − *z*_*m,t*_) maintains a sign such that:

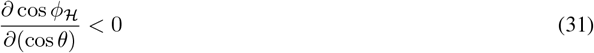

### Derivative of *ϕ*_*H*_ **with respect to** *θ*

We relate the change in angle *ϕ*_*H*_ to *θ* using the chain rule terms:

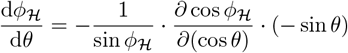

Substituting the derived terms:

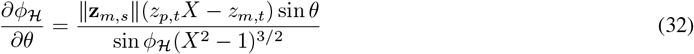

### Conclusion

For *θ* ∈ (0, *π*), we have sin *θ >* 0 and sin *ϕ*_*H*_ *>* 0. Under the standard geometric configuration where the ligand is distinct from the target, the numerator terms are strictly positive. Thus, we obtain:

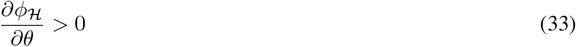

This confirms that as the tangent alignment angle *θ* increases (decreases), the hyperbolic exterior angle *ϕ*_ℋ_ strictly increases (decreases).

## Proof of Proposition 4.5

### Proposition A.7

*Let* **m**^*′*^ *be decomposed into tangential and radial components* 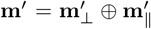. *Driven by the cone separation, if an active ligand is successfully entailed within the cone, i.e*. 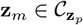, *then its tangent components satisfy the strict linear convergence constraint:*

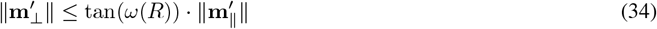

To prove Proposition 4.5, we first establish the following lemma regarding the relationship between the hyperbolic exterior angle and the Euclidean input angle defined below.

### Lemma A.8

*Due to the global convexity of geodesics in the negatively curved manifold, the hyperbolic exterior angle ϕ*_ℋ_ *strictly upper-bounds the Euclidean input angle ϕ*_Euc_ *for any finite, non-collinear interaction vector* **m**^*′*^ = **m** − **p**:

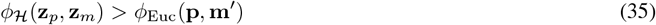

*Proof of Lemma A.8*. We provide a rigorous proof of the strict inequality *ϕ*_ℋ_ *> ϕ*_Euc_ for finite, non-collinear vectors 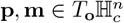.

### Convexity of Geodesic Pre-images

Since 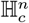 is a Hadamard manifold (simply connected with non-positive sectional curvature), it is a CAT(−*c*) space. A fundamental property of such spaces is that the squared distance function 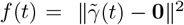 along any geodesic 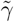 is strictly convex with respect to *t*, unless the geodesic passes through the origin.

Moreover, the radial metric factor in hyperbolic space, 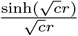, is strictly increasing with the radius *r*. Therefore, to minimize the hyperbolic path length, the geodesic *γ* must remain closer to the origin than the Euclidean straight line between **p** and **m**, implying that 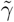 is strictly convex *towards the origin* relative to the Euclidean chord.

### Initial Tangent Vectors and Gauss Lemma

Let **m**^*′*^ = **m** − **p** be the Euclidean chord direction, and 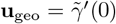 the initial tangent of the pre-image curve.

By the Gauss Lemma, the exponential map at the origin preserves angles between radial vectors and any tangent vector. Therefore, the angle between the outward radial direction **p** and **u**_geo_ in the tangent space equals the hyperbolic exterior angle *ϕ*_ℋ_ on the manifold, while the angle between **p** and **m**^*′*^ equals *ϕ*_Euc_.

### Convexity Implies Angle Amplification

Let *ψ*(**u**) denote the *interior angle* formed between −**p** (inward radial direction) and a vector **u**. Strict convexity of 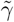 toward the origin implies

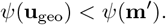

The exterior angle relative to the outward radial direction is defined as *ϕ* = *π* − *ψ*. Hence,

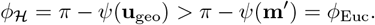

### Boundary Cases

Equality holds if and only if **p** and **m** are radially collinear (so that 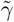 is a straight line through the origin). For vectors with very small magnitude or zero (which can be ignored), the same conclusion holds trivially as both angles approach zero.

### Conclusion

Thus, for any finite, non-collinear vectors **p, m**, the hyperbolic exterior angle strictly exceeds the Euclidean input angle:

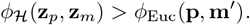

*Proof of Proposition 4.5*. Under the hierarchy assumption, if 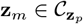, then the tangent components strictly satisfy:

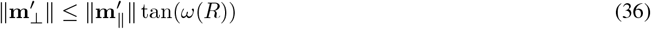

Combining the cone definition with Lemma A.8, we have the inequality chain: *ϕ*_Euclidean_ ≤ *ϕ*_ℋ_ ≤ *ω*(*R*). Since the tangent function is monotonically increasing on [0, *π/*2), it follows that tan(*ϕ*_Euclidean_) ≤ tan(*ω*(*R*)). In the Euclidean input space, the decomposition 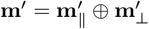 forms a right-angled triangle, where 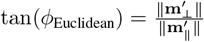. Substituting this identity into the inequality yields 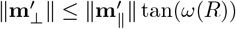.

## B. Analysis of Hyperbolic Margin Amplification via Euclidean Angular Separation

In the main text, we introduced the tangent separation loss ℒ_*tan*_ to enforce directional compatibility in the Euclidean tangent space. The motivation behind this design is that the hyperbolic geometry acts as a “geometric amplifier”: it converts angular separation in the tangent space into significant geodesic distance gaps on the manifold. In this section, we provide the theoretical justification for this strategy. We analyze the lower bound of the hyperbolic ranking margin—defined as the difference in geodesic distances 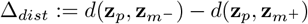 —and demonstrate that enforcing orthogonal separation in the Euclidean space guarantees a margin that scales linearly with the target radius.

### Proposition B.1

*Let R denote the norm of the target* **p**. *Let r*_+_ *and r*_−_ *denote the norms of the active ligand (***m**^+^*) and the decoy ligand (***m**^−^*), respectively. Assume the ligand embeddings are pushed towards the boundary of the manifold (r*_+_ ≫ 1,*r*_−_≫ 1*), a typical behavior in high-dimensional hyperbolic representations. Suppose the training objective successfully establishes the following geometric configurations:*

1. *The decoy is at least as far from the origin as the active ligand: r*_−_ ≥ *r*_+_ + Δ_*r*_.
2. *The active ligand is well-aligned with the target (small angle):* 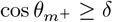.
3. *The decoy is geometrically isolated (large angle separation):* 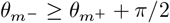.

*Then, the resulting hyperbolic ranking margin* Δ_*dist*_ *is lower-bounded by:*

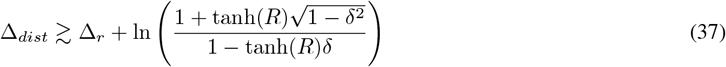

*Proof*. We analyze the geodesic distance *d*(**z**_*p*_, **z**_*m*_) using the Hyperbolic Law of Cosines:

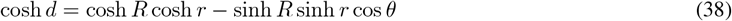

We assume that ligands are far from the origin (*r* ≫ 1). In this regime, we approximate cosh 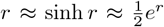. The distance equation simplifies to:

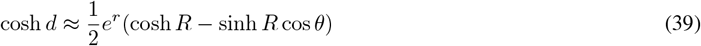

Using the approximation *d* ≈ ln(2 cosh *d*) for large distances, we substitute the expression above:

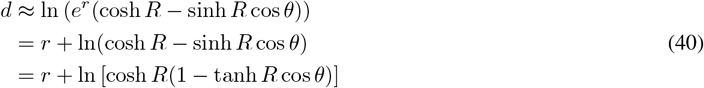

Notably, this derivation holds for any target radius *R*, as we did not approximate cosh *R*.

Now, we substitute this form into the definition of the ranking margin 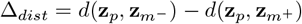. The ln(cosh *R*) term cancels out:

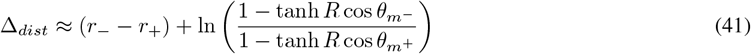

We now apply the three assumed geometric conditions to lower-bound this expression:

### Radial Contribution

From condition (1), *r*_−_ − *r*_+_ ≥ Δ_*r*_.

### Denominator Maximization (Active Alignment)

We maximize the denominator to find the lower bound. From condition (2), the worst-case active alignment is 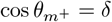:

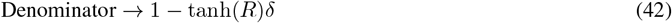

Denominator → 1 − tanh(*R*)*δ* (42)

### Numerator Minimization (Decoy Separation)

We minimize the numerator using condition (3): 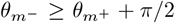. Considering the worst-case active angle 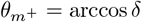, we have 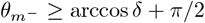. Since cosine is decreasing in [0, *π*], we bound 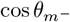:

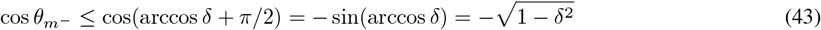

Substituting this upper bound into the numerator term yields:

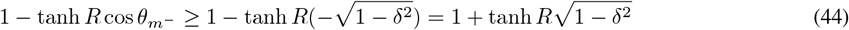

### Conclusion

Combining these terms yields the final inequality:

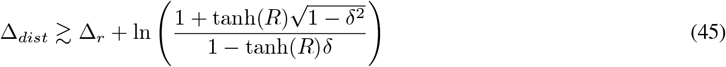

This completes the proof.

## C. Theoretical Guarantees of 4.3.1

### Problem Formulation

In the context of drug discovery, we aim to rank a set of ligands 𝒳 = {*x*_1_, …, *x*_*n*_} based on their binding affinity to a specific protein target. Let *f*_Θ_ : 𝒳 → ℝ denote a scoring function parameterized by Θ, where *s*_*i*_ = *f*_Θ_(*x*_*i*_) is the predicted affinity score.

#### Note on Notation

While the main text utilizes {*t*_1_, *t*_2_, …, *t*_*k*_} to denote ligands, for the sake of mathematical conciseness and to maintain consistency with standard ranking theory literature, we adopt the notation {*x*_1_, …, *x*_*n*_} throughout this Appendix. This shift is for notational simplicity and is made without loss of generality.

Unlike standard Information Retrieval (IR) tasks where relevance labels are discrete, biological affinity data is continuous but prone to experimental noise. To address this, we introduce high-confidence ranking lists to serve as the robust ground truth for this ranking formulation.

For a specific protein target, we employ a greedy algorithm to extract an ordered sequence of active ligands 𝒳_*active*_ from the raw assay data. To ensure ranking robustness, we enforce a significant affinity constraint: any adjacent ligands *x*_*i*_, *x*_*i*+1_ in the retained list must exhibit an affinity difference of at least 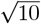-fold. This preprocessing converts continuous measurements into a rigorous ground truth ranking chain: for any pair with ranks *s < i*, the ligand *x*_*s*_ strictly dominates *x*_*i*_ (*x*_*s*_ *≻ x*_*i*_) with high confidence.

### Quadruplet Construction

Based on 𝒳_*active*_, we construct training instances as quadruplets *Q* = {*x*_1_, *x*_2_, *x*_3_, *x*_*decoy*_} through a two-step process:

1. **Exhaustive Active Enumeration:** We generate the set of all valid triplets { *x*_1_, *x*_2_, *x*_3_} ⊂ 𝒳_*active*_ such that their ranks satisfy *r*_1_ *< r*_2_ *< r*_3_. This exhaustive enumeration maximizes the utilization of pairwise ranking signals across all affinity gradients.
2. **Cluster-based Negative Sampling:** To pair each triplet with a high-quality decoy *x*_*decoy*_, we first cluster all training targets based on sequence similarity. For the current target, we randomly select a ligand associated with a target from a disjoint cluster. This cross-cluster strategy effectively precludes false negatives (randomly sampled binders), ensuring the decoy is biologically non-interacting.

Based on this construction, we formalize the existence of a deterministic ranking order:

#### Assumption C.1

For the constructed high-confidence ligand list 𝒳, there exists a unique ground truth permutation *y*^*^. For any pair of ranks *s < i*, the ligand at rank *s* satisfies a strict preference relation over the ligand at rank *i*:

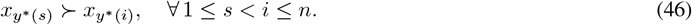

To optimize this ranking efficiently, we decompose the list into atomic units. We first define the local training objective used during stochastic optimization, and then the global objective it implicitly minimizes.

#### Definition C.2

Let 𝒯 = {(*s, i, k*) | 1 ≤ *s < i < k* ≤ *n*} be the set of all valid triplets consistent with *y*^*^. For any triplet *t* = (*s, i, k*) ∈ 𝒯, the atomic triplet loss is defined as a weighted combination of pairwise discrepancies using a convex, strictly decreasing surrogate function *ϕ*(·) (e.g., logistic loss):

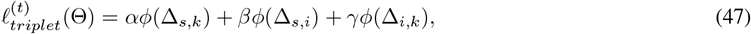

where 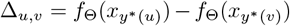 represents the score difference between rank *u* and *v*, and *α, β, γ* ≥ 0 are structural hyperparameters.

#### Definition C.3

We define the global ranking objective over the entire list 𝒳 as a distance-weighted summation of all pairwise losses:

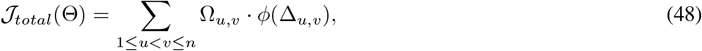

where the weighting matrix Ω ∈ ℝ^*n×n*^ is determined by the rank distance topology:

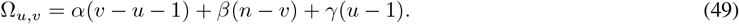

We now establish that our stochastic triplet sampling strategy is not merely a heuristic, but a mathematically exact method for optimizing the global objective.

#### Proposition C.4

*Assume a mini-batch contains the full set of triplets* 𝒯 *for a given ligand list* 𝒳. *The gradient of the aggregated atomic triplet losses with respect to the model parameters* Θ *is strictly identical to the gradient derived from the Global Weighted Pairwise Loss* 𝒥_*total*_. *That is:*

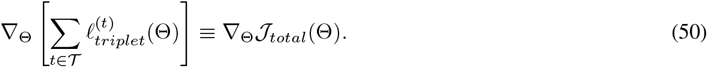

*Proof*. It suffices to prove the scalar algebraic identity 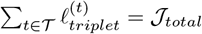. We determine the accumulated weight of an arbitrary pair (*u, v*) with *u* < *v* by enumerating all valid triplets (*s, i, k*) that contain this pair. The pair (*u, v*) appears in the summation in three mutually exclusive scenarios:

- **As the endpoints** (*s, k*): We set *s* = *u* and *k* = *v*. To form a valid triplet *s < i < k*, the intermediate index *i* can be any integer strictly between *u* and *v*. The number of such choices is (*v* − 1) − (*u* + 1) + 1 = *v* − *u* − 1. This term contributes with weight *α*.
- **As the first pair** (*s, i*): We set *s* = *u* and *i* = *v*. The third index *k* must satisfy *k > v* (since *k > i*). The number of valid choices for *k* is *n* − *v*. This term contributes with weight *β*.
- **As the second pair** (*i, k*): We set *i* = *u* and *k* = *v*. The first index *s* must satisfy *s < u* (since *s < i*). The number of valid choices for *s* is *u* − 1. This term contributes with weight *γ*.

Summing these counts, the total effective coefficient for the pair (*u, v*) is *α*(*v* − *u* − 1) + *β*(*n* − *v*) + *γ*(*u* − 1), which matches the definition of Ω_*u,v*_. Since the loss surfaces are identical, their gradients with respect to Θ are identical by the chain rule.

#### Remark C.5.

Although Proposition C.4 establishes the gradient equivalence between the local triplet objective and the global listwise objective, we explicitly adopt the triplet formulation rather than the full listwise approach for four critical reasons tailored to affinity ranking:

- **Magnitude-Aware Weighting:** Standard IR tasks typically operate on purely ordinal data (A is better than B), ignoring the scale of the difference. In contrast, our high-confidence affinity lists provide cardinal information: we know *how much* stronger a binder is. Our weighting matrix Ω (specifically, by setting *α* = 1 *> γ* = *β*) exploits this extra dimension: rank distance correlates with affinity magnitude. This naturally imposes heavier penalties on pairs with larger binding gaps, enforcing global structural consistency.
- **Data Efficiency:** Affinity datasets are significantly sparser than web-scale ranking logs. Decomposing lists into atomic triplets serves as a combinatorial data augmentation strategy, maximizing signal extraction from limited assay samples.
- **Implementation Flexibility:** Biological datasets are inherently jagged, with varying numbers of known actives per target. Listwise methods often require complex padding or truncation to handle variable sequence lengths. Triplet decomposition breaks these lists into atomic, fixed-size units, making the framework invariant to list length and significantly simplifying engineering implementation.
- **Robustness to Local Noise:** Wet-lab data inevitably contains noise. In global listwise objectives (e.g., Softmax-based losses), a single incorrect label, such as a weak binder mislabeled as strong, can propagate gradient errors across the entire list. In contrast, triplet optimization localizes these errors: invalid partial orders only affect specific atomic units, preserving the gradients of reliable pairs.

While Proposition C.4 guarantees optimization correctness, it remains to show that minimizing 𝒥_*total*_ effectively improves ranking quality. We define the Essential Loss to quantify discrete ranking violations.

To rigorous justify that our optimization objective aligns with standard ranking metrics, we first formally define the Normalized Discounted Cumulative Gain (Järvelin & Kekäläinen, 2002) and then prove that it is theoretically bounded by the Essential Loss defined below under a specific weighting scheme.

#### Definition C.6

Given the ground truth permutation *y*^*^ (Assumption C.1) and a scoring function *f*_Θ_ generating a predicted ranked list *π*_*f*_, the Normalized Discounted Cumulative Gain(NDCG) at cutoff *n* is defined as:

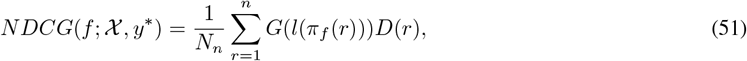

where *l*(·) is the relevance label derived from the affinity, *G*(·) is an increasing gain function (typically *G*(*z*) = 2^*z*^ − 1), *D*(·) is a decreasing position discount function (typically 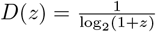), and *N*_*n*_ is the ideal normalization factor such that the max NDCG is 1.

We now connect this metric to the Essential Loss (Chen et al., 2009).

#### Definition C.7

Let *β*(*s*) be a rank-dependent importance weight. The Essential Loss *L*_*β*_ explicitly penalizes rank inversions:

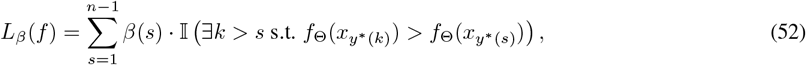

where 𝕀 (·) is the indicator function.

We specify the exact form of the rank-dependent weight *β*(*s*) required to establish a theoretical bound.

#### Definition C.8

To target NDCG optimization, we define the specific weight *β*_*NDCG*_(*s*) for the Essential Loss at each rank *s* as the product of the ground-truth gain and the position discount:

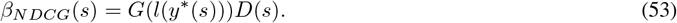

With this definition, we provide the fundamental theorem linking the discrete ranking error to our loss framework.

#### Theorem C.9

*For any scoring function f, the ranking error measured by* (1 − *NDCG*) *is strictly upper-bounded by the scaled Essential Loss* 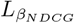. *Specifically:*

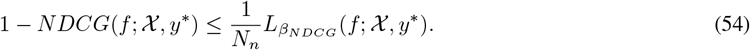

*Proof*. This result is adapted from Chen et al., 2009. We reformulate the NDCG summation indices from the predicted rank positions *r* to the ground truth positions *s*. Since *D*(*r*) is a strictly decreasing function, any rank inversion where an item *y*^*^(*s*) is ranked lower than *y*^*^(*k*) (with *k > s*) implies that *y*^*^(*s*) is pushed to a position with a smaller discount factor than ideal. Specifically, if an inversion occurs at *s*, the penalty contribution in the Essential Loss is weighted by *β*(*s*) = *G*(*l*(*y*^*^(*s*)))*D*(*s*). In contrast, the actual loss in NDCG is the difference in discounted gains, which is strictly upper-bounded by the full removal of the gain *G*(*l*(*y*^*^(*s*))) at position *s*. Thus, the weighted sum of inversion indicators (Essential Loss) dominates the metric error (1 − *NDCG*).

#### Theorem C.10

*The discrete Essential Loss is strictly upper-bounded by the differentiable Global Weighted Pairwise Loss, scaled by the geometric properties of the weighting matrix* Ω:

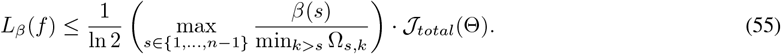

*Proof*. Let *E*_*s*_ = 𝕀 ( *∃k > s* : Δ_*s,k*_ *<* 0) denote the indicator of an inversion error at rank *s*. If *E*_*s*_ = 1, there exists at least one *k > s* such that Δ_*s,k*_ *<* 0. Since *ϕ*(*z*) = ln(1 + *e*^−*z*^) is strictly decreasing and *ϕ*(0) = ln 2, this implies *ϕ*(Δ_*s,k*_) *>* ln 2. The partial contribution of rank *s* to the global loss is 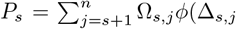). Due to the non-negativity of terms, *P*_*s*_ ≥ Ω_*s,k*_*ϕ*(Δ_*s,k*_) *>* (min_*j>s*_ Ω_*s,j*_) ln 2. Rearranging yields 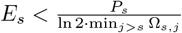. If *E*_*s*_ = 0, the inequality holds trivially since *P*_*s*_ ≥ 0. Summing over all ranks *s* weighted by *β*(*s*):

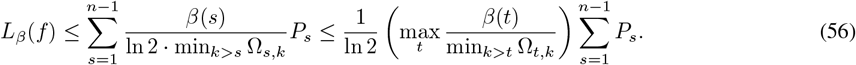

Substituting ∑ *P*_*s*_ = 𝒥_*total*_ completes the proof.

Finally, by chaining the bounds established in Theorem C.9 and Theorem C.10 (and its closed-form instantiation in Proposition C.12), we arrive at the fundamental consistency guarantee for our framework.

#### Corollary C.11

*Minimizing the Global Weighted Pairwise Loss 𝒥*_*total*_ *(and equivalently, the Triplet Loss) serves as a theoretically strictly bounded surrogate for maximizing the NDCG metric. Specifically, the ranking error is bounded by:*

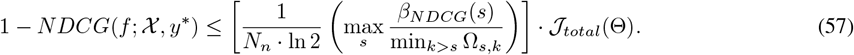

*This confirms that driving the training loss* 𝒥_*total*_ → 0 *strictly forces the ranking metric NDCG* → 1, *providing a rigorous theoretical foundation for the effectiveness of the triplet-based ranking loss*.

Before deriving the closed-form bound, we analyze the structural properties of the weighting matrix Ω to demonstrate the soundness of our loss design. Under the standard hyperparameter setting where *α* = 1 and *β* = *γ <* 1, the weight Ω_*s,k*_ for a pair (*s, k*) simplifies as follows:

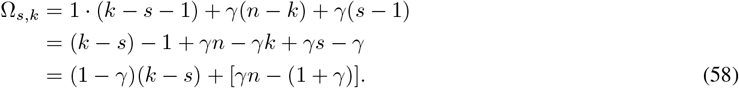

Here, (*k* − *s*) represents the rank distance between the two ligands in the ground truth. Since we assume *γ <* 1, the coefficient (1 − *γ*) is strictly positive. Consequently, Ω_*s,k*_ is **strictly monotonically increasing** with respect to the rank distance.

This monotonicity implies a crucial physical interpretation: the Global Weighted Pairwise Loss imposes significantly larger penalties on pairs that are far apart in the ground truth ranking. In our high-confidence affinity context, a large rank distance corresponds to a massive difference in binding affinity (e.g., a nanomolar binder vs. a micromolar non-binder). By assigning higher weights to these “easy-to-distinguish” but “costly-to-confuse” pairs, the objective function prioritizes the preservation of the global Structure-Activity Relationship (SAR) structure over minor local permutations, ensuring optimization soundness.

#### Proposition C.12

*Under the standard ranking weight* 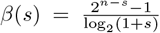 *and assuming structural hyperparameters α* = 1, *γ* = *β <* 1, *the bound simplifies to:*

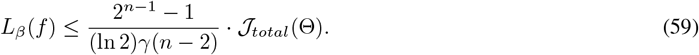

This proposition confirms that for high-confidence lists, our triplet-based optimization theoretically guarantees the suppression of rank inversions, providing a robust mechanism for affinity ranking.

## D. Details of Pairformer Features and Embedding Construction

To instantiate the geometric framework defined in the main text, we employ the Pairformer backbone Φ( ·) to extract comprehensive structural and interaction features. A critical implementation detail is the construction of a stable geometric baseline: we perform distinct forward passes to obtain the *reference state* (isolated protein) and the *bound state* (co-folded complex), which are then projected into the tangent space 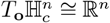 via domain-specific projection heads.

### 1. Dual-Pass Feature Extraction

Let 𝒫 denote the target protein sequence of length *N* and *L* denote the ligand SMILES string of length *M*.

- **Isolated Pass (Target Reference):** First, we feed only the protein sequence into the backbone to extract the ligand-agnostic target features. This ensures that the reference embedding remains constant across different ligands for the same target:

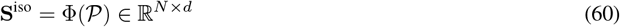

We aggregate this output to obtain the isolated protein single feature:

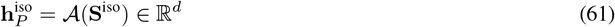
- **Co-folded Pass (Joint Complex):** Next, we feed the ptotein sequence and ligand SMILES into the backbone to simulate the binding complex. The model outputs the single representation **S**^cpx^ containing residue/atom-wise features and the pair representation **Z**^cpx^ capturing pairwise interactions:

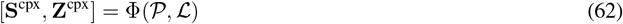

where **S**^cpx^ ∈ ℝ^(*N* +*M*)*×d*^ and **Z**^cpx^ ∈ ℝ^(*N* +*M*)*×*(*N* +*M*)*×c*^. From this pass, we extract three specific components: the ligand-modulated protein feature 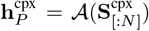, the ligand feature 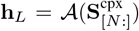, and the cross-interaction map 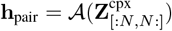.

### 2. Embedding Construction in Tangent Space

Based on the extracted components, we construct the final embeddings **e**_*x*_ (Target) and **e**_*y*_ (Ligand/Complex) for hyperbolic mapping.

- **Target Reference Vector (e**_*x*_): To establish a stable geometric anchor, the target embedding is encoded using solely the isolated protein feature via a Reference MLP. This ensures that the target’s position in the manifold serves as an invariant reference point:

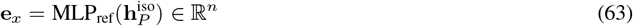
- **Joint Complex Vector (e**_*y*_): The complex embedding encapsulates the full binding context. We explicitly concatenate the ligand feature, the ligand-modulated protein feature, and the pairwise interaction feature, projecting them via a Complex MLP:

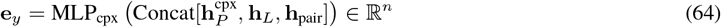

## E. Practical Implementation Details

### E.1 Architecture of Projection Networks

To map the raw embeddings from the pre-trained protein and ligand encoders into the hyperbolic tangent space, we employ two projection networks, *g*_*P*_ ( ·) and *g*_*L*_(·), as introduced in Section 4.2. While the input dimensions for the protein encoder (*d*_*p*_) and ligand encoder (*d*_*l*_) may differ, both networks share a unified architectural design aimed at satisfying the theoretical conditions outlined in Proposition 4.3.

Formally, each projector *g*(·) is implemented as a two-layer Multi-Layer Perceptron (MLP) with the following composition of operations:

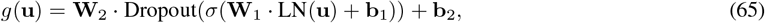

where:

- **u** denotes the input embedding (either **x** or **y**).
- LN(·) represents Layer Normalization, which stabilizes the input distribution.
- 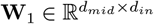 and **b**_1_ parameterize the first affine transformation.
- *σ*(·) is the ReLU activation function.
- 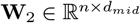 and **b**_2_ parameterize the final output layer, where *n* = 128 is the dimension of the tangent space.

### E.2 Theoretical Alignment

This specific architectural choice is designed to strictly adhere to the assumptions required for breaking the Euclidean Lipschitz barrier:

- **Lipschitz Continuity:** Since LayerNorm, Linear transformations, and ReLU activations are all Lipschitz continuous functions, their composition *g*(·) remains Lipschitz continuous, satisfying the precondition of Proposition 4.3.
- **Unboundedness:** Crucially, the final layer is a pure linear transformation (parameterized by **W**_2_, **b**_2_) without any bounding activation functions (e.g., Tanh or Sigmoid) or normalization. This ensures that the image of the projector is unbounded (*g*(ℝ^*d*^) = ℝ^*n*^), satisfying Assumption 1 of Proposition 4.3.
- **Non-vanishing Gradient:** The use of ReLU and standard weight initialization ensures that the Jacobian maintains full rank almost everywhere, preventing gradient collapse and satisfying Assumption 2.

### E.3 Hyperparameter Settings

We detail the specific hyperparameter configurations used in our experiments below.

- **Data Tuple Configuration:** We construct training instances as tuples of size *k* = 3.
- **Proximity-Aware Weighting:** For the term *ψ*_*ij*_, *α* = 2 and *σ* = 100.
- **Ranking Loss Coefficients:** The pairwise weights in the ranking objective ℒ_*rank*_, are set to *λ*_13_ = 1.0, *λ*_12_ = 0.7, and *λ*_23_ = 0.7.
- **Geometric Loss Coefficients:** In the geometric regularization term ℒ_*geo*_, *λ*_*tan*_ = 1.5, and *λ*_*cone*_ = 0.01.

### E.4. Training Protocol

We optimize the model parameters using the AdamW optimizer (Loshchilov & Hutter, 2017). We set the weight decay coefficient to 1 *×* 10^−3^ to prevent overfitting. For the learning rate schedule, we employ a linear warmup strategy to stabilize early training dynamics: the learning rate is linearly increased from 0 to a peak value of 2 *×* 10^−4^ over the first 200 warmup steps.

## F. Extended Case Studies

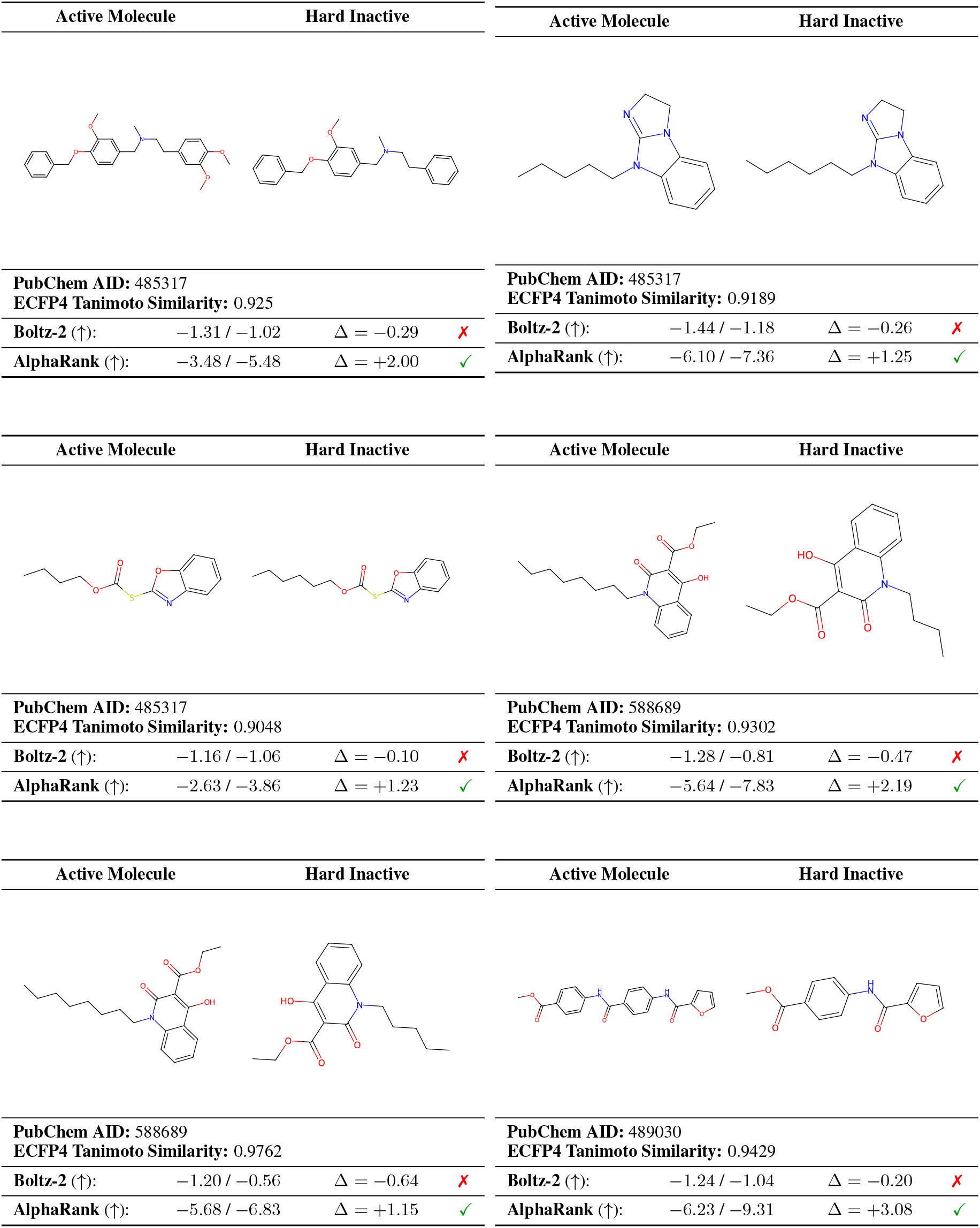

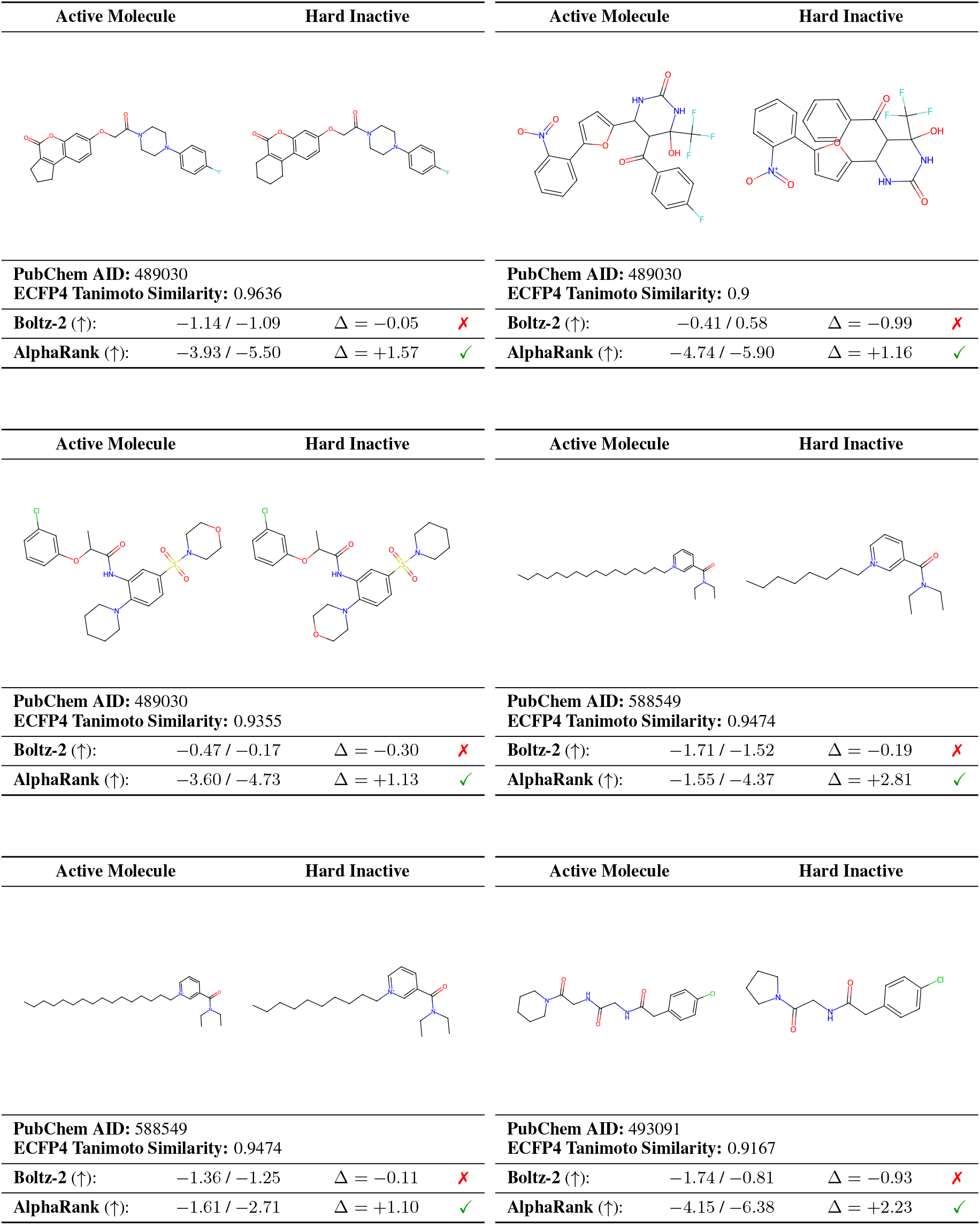

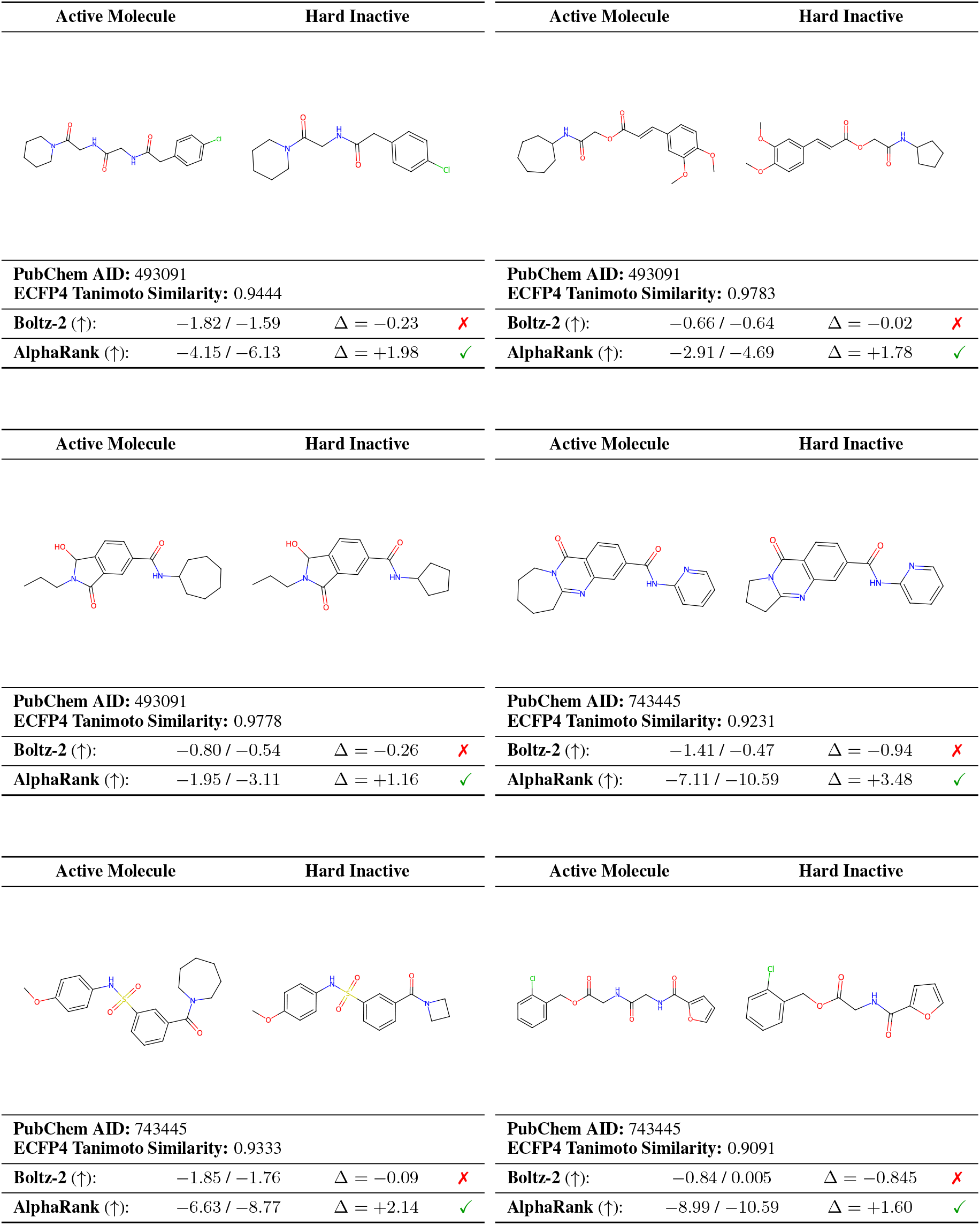

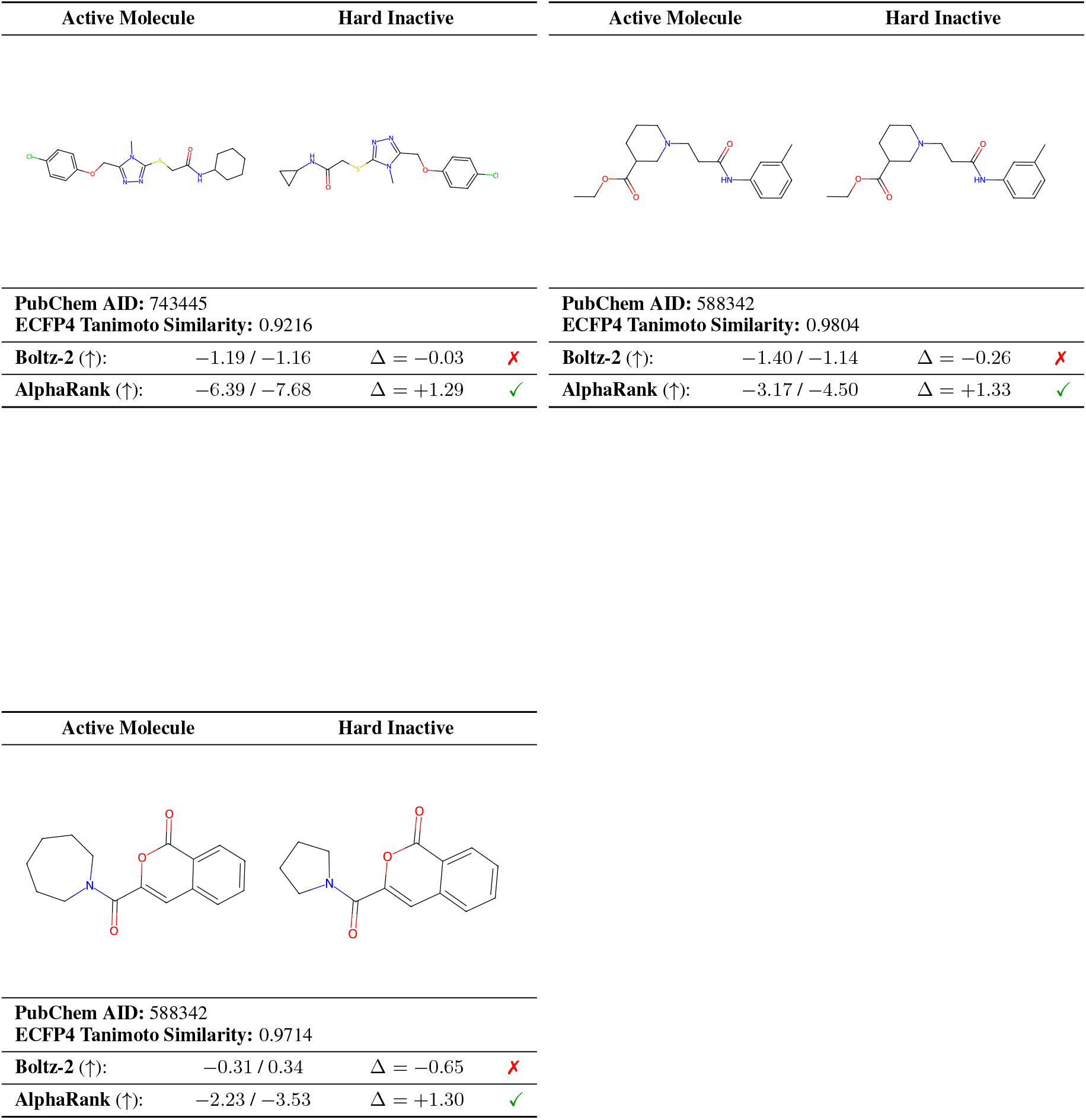

## References

Chamberlain, B. P., Clough, J., and Deisenroth, M. P. Neural embeddings of graphs in hyperbolic space. arXiv preprint 1705.10359, 2017.

Chen, W., Liu, T.-Y., Lan, Y., Ma, Z.-M., and Li, H. Ranking measures and loss functions in learning to rank. Advances in Neural Information Processing Systems, 22, 2009.

Chothia, C. and Janin, J. Principles of protein–protein recognition. Nature, 256(5520):705–708, 1975.

Desai, K., Nickel, M., Rajpurohit, T., Johnson, J., and Vedantam, S. R. Hyperbolic image-text representations. In International Conference on Machine Learning, pp. 7694–7731. PMLR, 2023.

Feng, B., Liu, Z., Yang, M., Zou, J., Cao, H., Li, Y., Zhang, L., and Wang, S. A foundation model for protein-ligand affinity prediction through jointly optimizing virtual screening and hit-to-lead optimization. bioRxiv, pp. 2025–02, 2025.

Feng, S., Li, M., Jia, Y., Ma, W.-Y., and Lan, Y. Proteinligand binding representation learning from fine-grained interactions. In The Twelfth International Conference on Learning Representations, 2024.

Friesner, R. A., Banks, J. L., Murphy, R. B., Halgren, T. A., Klicic, J. J., Mainz, D. T., Repasky, M. P., Knoll, E. H., Shelley, M., Perry, J. K., et al. Glide: a new approach for rapid, accurate docking and scoring. 1. method and assessment of docking accuracy. Journal of medicinal chemistry, 47(7):1739–1749, 2004.

Ganea, O., Bécigneul, G., and Hofmann, T. Hyperbolic entailment cones for learning hierarchical embeddings. In International conference on machine learning, pp. 1646–1655. PMLR, 2018.

Genheden, S. and Ryde, U. The mm/pbsa and mm/gbsa methods to estimate ligand-binding affinities. Expert opinion on drug discovery, 10(5):449–461, 2015.

Huang, K., Fu, T., Glass, L. M., Zitnik, M., Xiao, C., and Sun, J. Deeppurpose: a deep learning library for drug– target interaction prediction. Bioinformatics, 36(22-23): 5545–5547, 2020.

Järvelin, K. and Kekäläinen, J. Cumulated gain-based evaluation of ir techniques. ACM Transactions on Information Systems (TOIS), 20(4):422–446, 2002.

Jumper, J., Evans, R., Pritzel, A., Green, T., Figurnov, M., Ronneberger, O., Tunyasuvunakool, K., Bates, R., Žídek, A., Potapenko, A., et al. Highly accurate protein structure prediction with alphafold. nature, 596(7873):583–589, 2021.

Kong, X., Huang, W., and Liu, Y. Generalist equivariant transformer towards 3d molecular interaction learning. In Forty-first International Conference on Machine Learning, 2024.

Lam, H. Y. I., Guan, J. S., Ong, X. E., Pincket, R., and Mu, Y. Protein language models are performant in structurefree virtual screening. Briefings in Bioinformatics, 25(6):bbae480, 2024.

Le, M., Roller, S., Papaxanthos, L., Kiela, D., and Nickel, M. Inferring concept hierarchies from text corpora via hyperbolic embeddings. In Proceedings of the 57th annual meeting of the association for computational linguistics, pp. 3231–3241, 2019.

Loshchilov, I. and Hutter, F. Decoupled weight decay regularization. arXiv preprint 1711.05101, 2017.

Mobley, D. L. and Dill, K. A. Binding of small-molecule ligands to proteins:”what you see” is not always “what you get”. Structure, 17(4):489–498, 2009.

Nickel, M. and Kiela, D. Poincaré embeddings for learning hierarchical representations. Advances in neural information processing systems, 30, 2017.

Öztürk, H., Özgür, A., and Ozkirimli, E. Deepdta: deep drug–target binding affinity prediction. Bioinformatics, 34(17):i821–i829, 2018.

Pal, A., van Spengler, M., di Melendugno, G. M. D., Flaborea, A., Galasso, F., and Mettes, P. Compositional entailment learning for hyperbolic vision-language models. arXiv preprint 2410.06912, 2024.

Passaro, S., Corso, G., Wohlwend, J., Reveiz, M., Thaler, S., Somnath, V. R., Getz, N., Portnoi, T., Roy, J., Stark, H., et al. Boltz-2: Towards accurate and efficient binding affinity prediction. BioRxiv, 2025.

Schindler, C. E., Baumann, H., Blum, A., Bose, D., Buchstaller, H.-P., Burgdorf, L., Cappel, D., Chekler, E., Czodrowski, P., Dorsch, D., et al. Large-scale assessment of binding free energy calculations in active drug discovery projects. Journal of Chemical Information and Modeling, 60(11):5457–5474, 2020.

Team, B. A. A., Chen, X., Zhang, Y., Lu, C., Ma, W., Guan, J., Gong, C., Yang, J., Zhang, H., Zhang, K., et al. Protenix-advancing structure prediction through a comprehensive alphafold3 reproduction. bioRxiv, pp. 2025–01, 2025.

Trott, O. and Olson, A. J. Autodock vina: improving the speed and accuracy of docking with a new scoring function, efficient optimization, and multithreading. Journal of computational chemistry, 31(2):455–461, 2010.

Veith, H., Southall, N., Huang, R., James, T., Fayne, D., Artemenko, N., Shen, M., Inglese, J., Austin, C. P., Lloyd, D. G., et al. Comprehensive characterization of cytochrome p450 isozyme selectivity across chemical libraries. Nature biotechnology, 27(11):1050–1055, 2009.

Vendrov, I., Kiros, R., Fidler, S., and Urtasun, R. Orderembeddings of images and language. arXiv preprint 1511.06361, 2015.

Wang, J., Zhu, W., Gao, B., Hong, X., Zhang, Y.-Q., Ma, W.-Y., and Lan, Y. Learning protein-ligand binding in hyperbolic space. arXiv preprint 2508.15480, 2025.

Wang, L., Wu, Y., Deng, Y., Kim, B., Pierce, L., Krilov, G., Lupyan, D., Robinson, S., Dahlgren, M. K., Greenwood, J., et al. Accurate and reliable prediction of relative ligand binding potency in prospective drug discovery by way of a modern free-energy calculation protocol and force field. Journal of the American Chemical Society, 137(7):2695–2703, 2015.

Yang, Z., Zhong, W., Lv, Q., Dong, T., Chen, G., and Chen, C. Y.-C. Interaction-based inductive bias in graph neural networks: enhancing protein-ligand binding affinity predictions from 3d structures. IEEE Transactions on Pattern Analysis and Machine Intelligence, 2024.

Yu, J., Li, Z., Chen, G., Kong, X., Hu, J., Wang, D., Cao, D., Li, Y., Huo, R., Wang, G., et al. Computing the relative binding affinity of ligands based on a pairwise binding comparison network. Nature Computational Science, 3 (10):860–872, 2023.

Zdrazil, B., Felix, E., Hunter, F., Manners, E. J., Blackshaw, J., Corbett, S., De Veij, M., Ioannidis, H., Lopez, D. M., Mosquera, J. F., et al. The chembl database in 2023: a drug discovery platform spanning multiple bioactivity data types and time periods. Nucleic acids research, 52 (D1):D1180–D1192, 2024.

Zwanzig, R. W. High-temperature equation of state by a perturbation method. i. nonpolar gases. The Journal of Chemical Physics, 22(8):1420–1426, 1954.

